# Brain region-specific changes in neurons and glia and dysregulation of dopamine signaling in *Grin2a* mutant mice

**DOI:** 10.1101/2022.11.15.516665

**Authors:** Zohreh Farsi, Ally Nicolella, Sean K Simmons, Sameer Aryal, Nate Shepard, Kira Brenner, Sherry Lin, Linnea Herzog, Wangyong Shin, Vahid Gazestani, Bryan Song, Kevin Bonanno, Hasmik Keshishian, Steven A Carr, Evan Macosko, Sandeep Robert Datta, Borislav Dejanovic, Eunjoon Kim, Joshua Z Levin, Morgan Sheng

## Abstract

Schizophrenia disease mechanisms remain poorly understood, in large part due to a lack of valid animal models. Rare heterozygous loss-of-function mutations in *GRIN2A*, encoding a subunit of the NMDA (N-methyl-d-aspartate) receptor, greatly increase the risk of schizophrenia. By transcriptomic, proteomic, electroencephalogram (EEG) recording and behavioral analysis, we report that heterozygous *Grin2a* mutant mice show: (i) large-scale gene expression changes across multiple brain regions and in neuronal (excitatory and inhibitory) and non-neuronal cells (astrocytes, oligodendrocytes); (ii) evidence of reduced activity in prefrontal cortex and increased activity in hippocampus and striatum; (iii) elevated dopamine signaling in striatum; (iv) altered cholesterol biosynthesis in astrocytes; (v) reduction of glutamatergic receptor signaling proteins in the synapse; (iv) heightened gamma oscillation power in EEG; (vi) aberrant locomotor behavioral pattern opposite of that induced by antipsychotic drugs. These findings reveal potential pathophysiologic mechanisms, provide support for both the “hypo-glutamate” and “hyper-dopamine” hypotheses of schizophrenia, and underscore the utility of *Grin2a*-deficient mice as a new genetic model of schizophrenia.

## INTRODUCTION

Schizophrenia (SCZ) is a prevalent and disabling mental illness. Many decades after its recognition as a disease, the pathophysiologic mechanisms underlying SCZ remain mostly unknown. Existing pharmacological treatments for SCZ arose from serendipity, have limited efficacy and major side effects, underscoring the need for deeper understanding of SCZ disease mechanism to enable discovery of new therapies. Understanding of SCZ mechanism has been severely hampered by a lack of valid animal models.

SCZ has genetic and environmental risk factors, with an estimated heritability of 60-80% (Owen et al., 2016; Sullivan et al., 2003). Many common genetic variants have been identified by genome-wide association study (GWAS) of SCZ, including 287 in a recent analysis of >300,000 individuals most of which have small effects on disease risk (odds ratio (OR) usually less than 1.1) (Trubetskoy et al., 2022). Recent exome sequencing studies have uncovered rare genetic variants associated with SCZ, including protein truncating variants (PTVs) (Genovese et al., 2016; Singh et al., 2022; Singh et al., 2017) that have large effects on disease risk (OR = 2 - 60) (Singh et al., 2022). The most recent Schizophrenia Exome Sequencing Meta-Analysis (SCHEMA; ∼25,000 SCZ cases, ∼100,000 controls) has identified ten high-confidence genes (hereafter referred as SCHEMA genes) with loss-of-function (LoF) mutations associated with SCZ at exome-wide significance (Singh et al., 2022), including genes encoding synaptic signaling proteins (*TRIO*) and postsynaptic glutamate receptors (*GRIN2A* and *GRIA3*). These SCHEMA variants appear to be predominantly LoF mutations, suggesting that heterozygous loss of these genes is sufficient to confer substantial risk of SCZ (Rivas et al., 2015; Singh et al., 2022).

SCHEMA gene LoF mutations can be introduced in mice to create disease models with *bona fide* human-genetic validity, unlike many previous mouse genetic models of SCZ in which the mutated candidate genes have questionable links to human schizophrenia (see for example, (Sullivan, 2013)). Characterization of SCHEMA mouse mutants by comprehensive approaches can not only uncover potential disease mechanisms underlying SCZ, but also reveal phenotypes at the molecular, cellular, and network levels that can be compared with human patient data to discover SCZ disease biomarkers.

*GRIN2A*, encoding the GluN2A subunit of the NMDA receptor (NMDAR), is one of the ten exome-wide-significant SCHEMA genes (Singh et al., 2022); additionally, there is strong support for *GRIN2A* as a SCZ risk gene from GWAS fine-mapping (Trubetskoy et al., 2022). Besides SCZ, *GRIN2A* has been associated with neurodevelopmental disorders as well as epilepsy and speech disorders (Endele et al., 2010; Lesca et al., 2013; Pierson et al., 2014; Strehlow et al., 2019).

However, the *GRIN2A* association with SCZ is largely driven by PTVs (Singh et al., 2022) which are likely LoF variants (Rivas et al., 2015), while associations with neurodevelopmental disorder and epilepsy are predominantly through missense mutations that are clustered in the transmembrane and linker domains of *GRIN2A*, suggesting an alternate or gain-of-function mechanism (Chen et al., 2017; Strehlow et al., 2019; Yuan et al., 2014).

NMDAR hypofunction has long been hypothesized as a mechanism underlying SCZ pathophysiology, partly because NMDA receptor antagonists such as phencyclidine or ketamine at low concentrations can induce SCZ-like symptoms in healthy individuals (Moghaddam and Javitt, 2012). The discovery of *GRIN2A* as a LoF SCZ risk gene provides indirect human genetics support for the hypo-NMDAR function hypothesis of SCZ. In this context, it is notable that *Grin2a* expression emerges only postnatally in juvenile rodents (Sheng et al., 1994), reminiscent of SCZ onset that occurs typically in adolescence and young adulthood. Finally, NMDARs play crucial roles in excitatory synaptic transmission and plasticity (Paoletti et al., 2013), and synaptic dysfunction is implicated in SCZ by human genetics (Hall and Bray, 2022; Hall et al., 2015; Trubetskoy et al., 2022) and by postmortem gene expression studies (Gandal et al., 2018; Ruzicka et al., 2021).

To gain mechanistic insights into the role of *GRIN2A* in the pathophysiology of SCZ, we performed genome-wide mRNA expression profiling of multiple brain regions in *Grin2a* heterozygous and homozygous knockout mice at several ages, as well as proteomic analysis of purified synapses. We chose to study an existing *Grin2a* knockout mouse line (Kadotani et al., 1996) because it is representative of multiple SCHEMA protein truncating variants and damaging missense mutations in the *GRIN2A* gene which are predicted to be LoF mutations (Rivas et al., 2015; Singh et al., 2022). For additional insights, we compared the brain transcriptomic profile of *Grin2a* heterozygous mutants with that of *Grin2b* heterozygous knock-in mice carrying a null mutation (C456Y) that is associated with autism spectrum disorder (ASD) (Shin et al., 2020). *GRIN2B* encodes the GluN2B subunit of the NMDAR, which is expressed from embryonic stages of brain development (Sheng et al., 1994), and has so far not been linked to SCZ.

These comprehensive multi-omics investigations of *Grin2a* mutant mice--performed in the context of EEG studies showing abnormalities in brain oscillations, and automated behavioral analysis (MoSeq) showing locomotor perturbations--revealed unexpected molecular pathway changes (e.g. cholesterol biosynthesis in astrocytes) and systems disturbances (e.g. hyper-dopaminergic state in striatum). Together, our data builds a molecular-to-systems level picture of *Grin2a* heterozygous mutant mice as a compelling genetic animal model of SCZ and as a resource for deeper understanding of SCZ disease mechanism and therapeutics discovery.

## RESULTS

### Multiple brain regions across development are differentially affected in *Grin2a* and *Grin2b* mutant mice

To capture a comprehensive picture of the brain transcriptome in *Grin2a* and *Grin2b* mutant mice across different ages, we first performed bulk RNA-seq analysis on *Grin2a*^+/-^, *Grin2a*^-/-^, *Grin2b*^+/C456Y^ mice (*Grin2b* ^C456Y /C456Y^ is lethal), and their wild-type littermates at 2, 4, 12, and 20 weeks of age (n = 4-5 per genotype per timepoint). Multiple brain regions were examined including prefrontal cortex (PFC), hippocampus, somatosensory cortex (SSC), striatum, and thalamus (Fig. 1A). The same brain regions from different animals clustered together in the principal component analysis (PCA), indicating consistency of tissue dissection (Fig. S1A).

**Figure 1.**
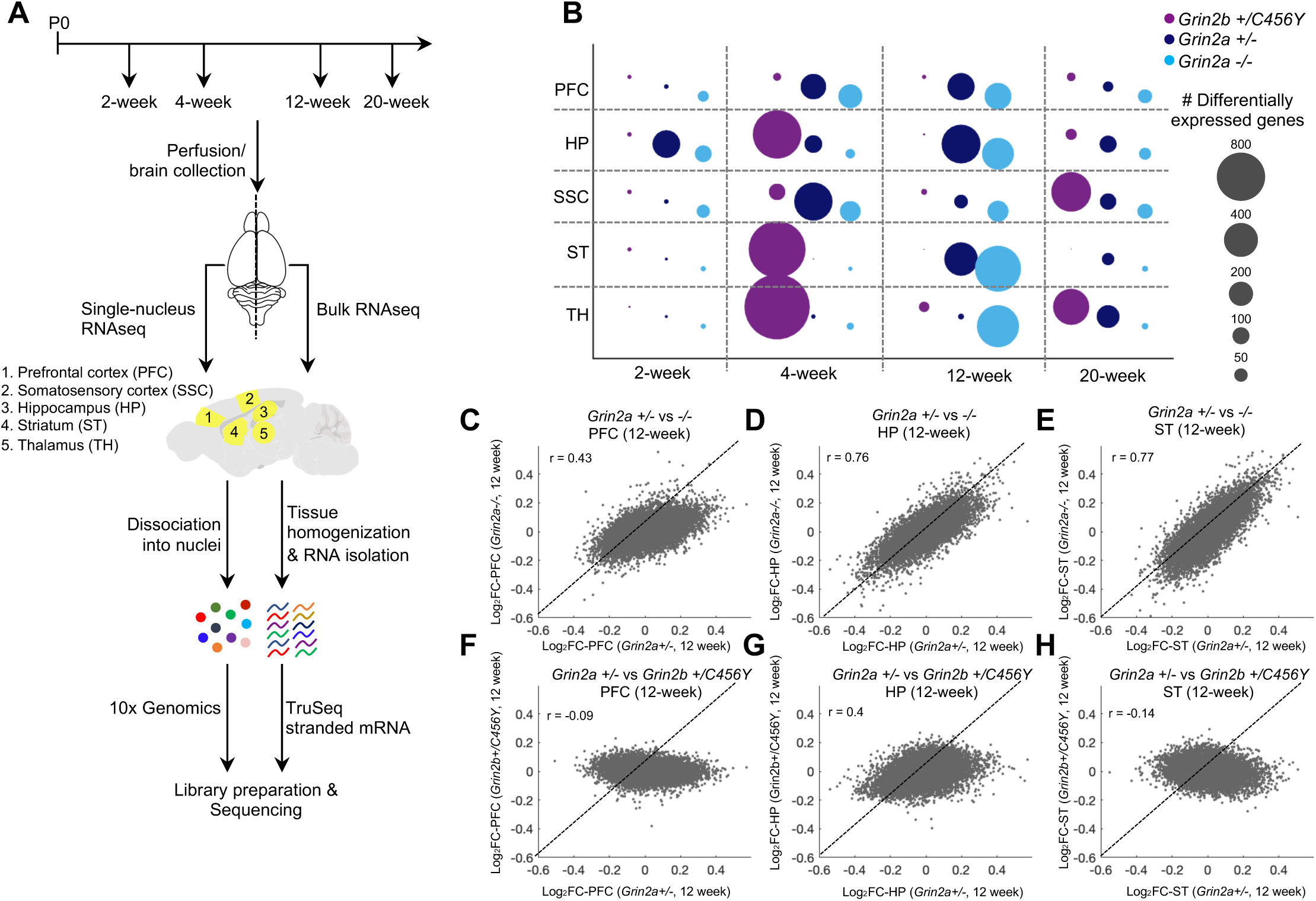
*Grin2a* and *Grin2b* mutant mice show widespread transcriptomic changes in different brain regions. (A) Schematic diagram of bulk and snRNA-seq characterization of multiple brain regions in *Grin2a* and *Grin2b* mutant mice at four ages. (B) Bubble plot showing the number of DEGs (FDR < 0.05, absolute value of Log2FC > 0.2) in each brain region and age in *Grin2a* and *Grin2b* mutants. (**C-H**) Gene expression correlation for *Grin2a^+/-^* versus *Grin2a^-/-^* (**C-E**) and in *Grin2a^+/-^* versus *Grin2b^+/C456Y^* at 12 weeks (**F-H**) in the prefrontal cortex (**C, F**), hippocampus (**D, G**), and striatum (**E, H**). In **C-H**, pearson’s r correlation values are indicated on the plots..

By bulk RNA-seq, a number of differentially expressed genes (DEGs, defined as genes with changes in mRNA expression versus wild types with false discovery rate (FDR) < 0.05 and absolute value of Log2Fold Change (Log2FC) > 0.2) were identified in both *Grin2a* and *Grin2b* mutants in all five brain regions and at all four ages investigated (Fig. 1B; see Tables S1-2 for all DEGs across all datasets). We validated a subset of these DEGs using qRT-PCR, showing the overall robustness of the RNA-seq results (Fig. S1B). *Grin2b*^+/C456Y^ showed the most DEGs at 4 weeks (total of 3,611 DEGs across all tested brain regions), as compared to the other tested ages (Fig. 1B). In *Grin2a* mutants, however, the greatest number of DEGs was found at 12 weeks (1,287 in *Grin2a*^+/-^ and 2,215 in *Grin2a*^-/-^, across tested brain regions) (Fig. 1B). Comparing the transcriptomic changes of all individual genes in the same brain region at 12 weeks versus 4 weeks, we noted larger Log2Fold changes at 12 weeks than at 4 weeks in most brain regions in the *Grin2a^+/-^* mutants (Fig. S1C-G) while in *Grin2b^+/C456Y^* mutants larger Log2Fold changes occurred at 4 weeks of age (Fig. S1H-L). There were relatively few DEGs for *Grin2a* or *Grin2b* mutants at 2 weeks. We therefore concentrated our subsequent analyses on 12 weeks for *Grin2a* and 4 weeks for *Grin2b* mutants.

Focusing on differential brain-region effects, we compared the heterozygous mutants *Grin2b*^+/C456Y^ (4 weeks) versus *Grin2a*^+/-^ (12 weeks). The hippocampus and striatum showed large transcriptomic changes in both genotypes. The PFC, which is associated with cognitive impairment in SCZ (Sakurai et al., 2015), showed more DEGs in *Grin2a*^+/-^ than in *Grin2b*^+/C456Y^ (Fig. 1B) whereas the thalamus showed a larger number of DEGs in *Grin2b*^+/C456Y^, even though the number of samples and the average sequencing depth per brain region were similar between the two genotypes (see Table S3). These results suggest that brain regions and circuitries might be differentially affected by *Grin2a* and *Grin2b* LoF.

Comparing heterozygous versus homozygous *Grin2a* mutant mice, it was notable that *Grin2a*^+/-^ showed comparable numbers of DEGs to *Grin2a*^-/-^ in most brain regions (see for example, PFC and hippocampus) at all four ages (Fig. 1B), suggesting that loss of just one copy of *Grin2a* is sufficient to exert profound transcriptomic changes across the brain.

A transcriptome-wide comparison showed that mRNA changes of individual genes (Log2FC values) were well correlated between *Grin2a*^+/-^ and *Grin2a*^-/-^ mutants in all brain regions studied (Pearson’s r = 0.43 - 0.77 (see Methods); Fig. 1C-E, Fig. S1M, N). These data demonstrate that the transcriptome profiles of the same brain regions are considerably overlapping in *Grin2a^+/-^* and *Grin2a^-/-^* mutants, and that the heterozygous *Grin2a* LoF has as much effect than homozygous *Grin2a* LoF on the overall transcriptome. By contrast, the transcriptome profile of *Grin2a*^+/-^ was poorly or even anti-correlated with that of *Grin2b*^+/C456Y^ in the same brain regions (Pearson’s r < 0.1; Fig. 1F, H, Fig. S1O, P), except for hippocampus (Pearson’s r = 0.4; Fig. 1G). Thus, loss of one copy of *Grin2a* or *Grin2b* causes markedly different global transcriptomic changes at the same age, despite both genes encoding subunits of the NMDARs.

### Differential alteration of diverse molecular pathways in brain regions of *Grin2a* and *Grin2b* mutants

To gain insight into the biological processes that were altered in *Grin2a* and *Grin2b* mutant mice, we performed Gene Set Enrichment Analysis (GSEA) (Korotkevich et al., 2021; Subramanian et al., 2005) (see Methods) on bulk RNA-seq data in all brain regions, ages, and genotypes (see Tables S4-5 for all significant GO terms across all datasets).

In the GSEA analysis of 4-week *Grin2b*^+/C456Y^ and 12-week *Grin2a* mutants, a number of pathways (annotated by Gene Ontology (GO) terms) were found to be significantly (FDR < 0.05) enriched among up- and down-regulated genes (hereafter referred to as upregulated or downregulated pathways/GO terms, respectively) across multiple brain regions in both *Grin2a* and *Grin2b* mutant mice. Notably, these ‘convergent pathways’ were related to synapses, ribosomes, oxidative phosphorylation (OXPHOS), neuron development and RNA processing/splicing (Fig. 2A). In *Grin2a*^+/-^ and *Grin2a*^-/-^ mice, synaptic GO terms including glutamatergic and *gamma*-aminobutyric acid (GABA)ergic synapses were downregulated in the PFC and hippocampus, but the same GO terms were upregulated in the striatum (Fig. 2A). Consistent with the anti-correlated pathway change, many individual genes within the synapse GO term showed negative correlation in the PFC versus the striatum in *Grin2a^+/-^* mice (Pearson’s r = -0.23; Fig. S1Q). This opposite directionality of change in the PFC and hippocampus versus the striatum was also observed in *Grin2a* mutant mice for GO terms related to neuron development (down in PFC and hippocampus; up in striatum), and for OXPHOS and ribosomal GO terms (up in PFC and hippocampus; down in striatum) (Fig. 2A, Fig. S1R, S). By contrast, in *Grin2b*^+/C456Y^ mice, OXPHOS and ribosomal GO terms were significantly upregulated in all tested brain regions (Fig. 2A). Of particular note, in most studied brain regions in *Grin2a* and *Grin2b* mutants, the OXPHOS and ribosomal GO terms changed together in the same direction, but in the opposite direction with respect to synaptic GO terms.

**Figure 2.**
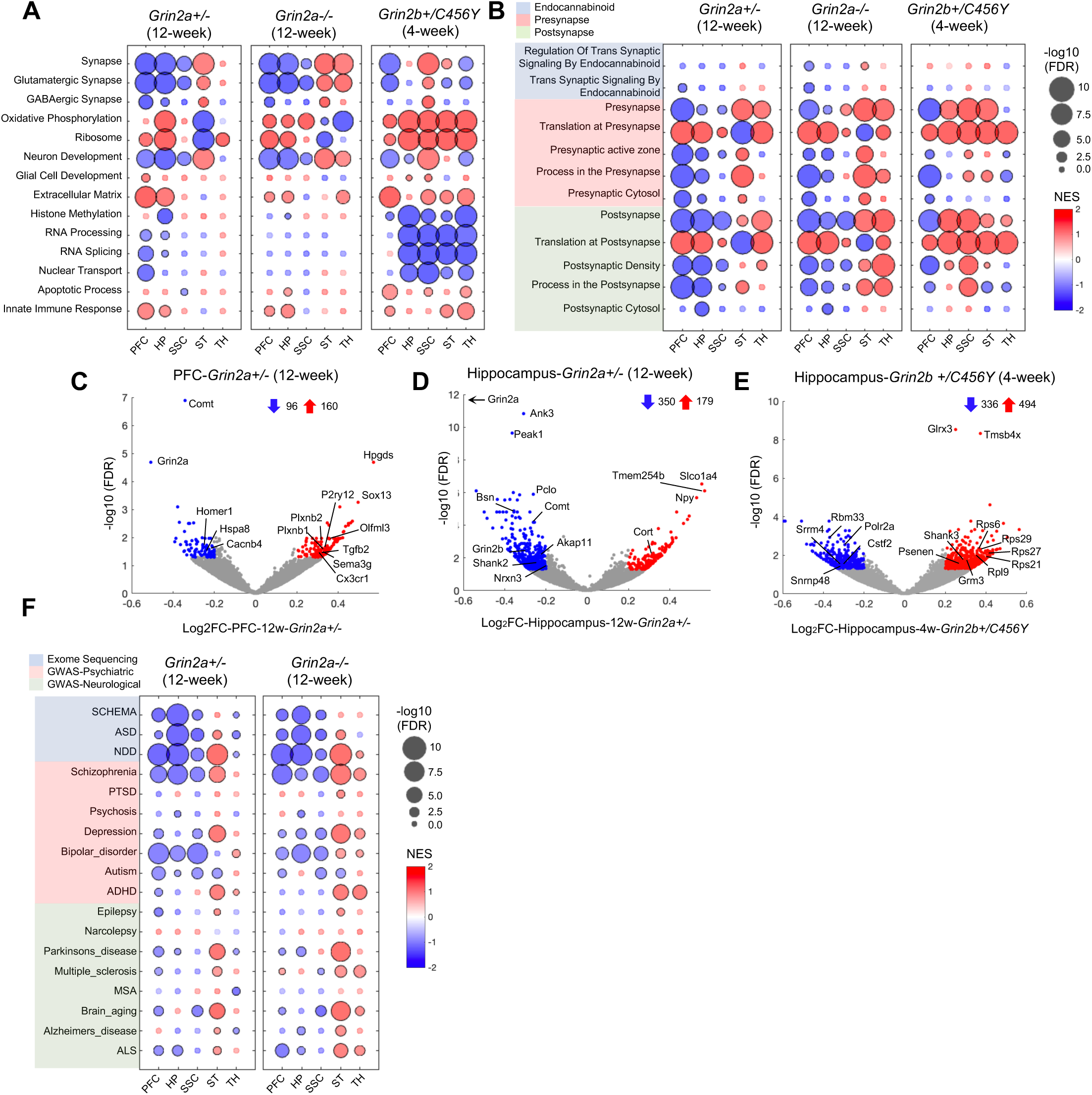
GSEA reveals significant changes in multiple molecular pathways in *Grin2a* and *Grin2b* mutant mice. (**A-B**) GSEA results for the studied brain regions in *Grin2a* (12-week), and *Grin2b* (4-week) mutants showing enrichment pattern of a selection of GO terms from MSigDB (**A**) and SynGO defined synaptic ontology terms (**B**). (**C-E**) Volcano plots showing all significant DEGs (blue and red circles represent down- and upregulated DEGs, respectively) in the prefrontal cortex (**C**) and hippocampus (**D**) of 12-week *Grin2a^+/-^* as well as hippocampus of 4-week *Grin2b^+/C456Y^* (**E**). Numbers next to blue and red arrows indicate the number of down- and upregulated DEGs, respectively. (**F**) GSEA results for the studied brain regions in *Grin2a* mutants (12-week) showing enrichment pattern of risk genes from psychiatric and neurological diseases as identified by exome sequencing and GWAS studies. In **A**, **B** and **F** circles with black outlines indicate significance (FDR < 0.05). In **D**, black arrow indicates that *Grin2a* is out of range of the plot. In **C**-**E** genes of interest are labeled on volcano plots. PFC, prefrontal cortex; HP, hippocampus; TH, thalamus; SSC, somatosensory cortex; ST, striatum; SCHEMA, schizophrenia exome sequencing meta-analysis; ASD; autism spectrum disorder; NDD; neurodevelopmental disorder; PTSD, post-traumatic stress disorder; ADHD, attention deficit hyperactivity disorder; MSA, multiple system atrophy; ALS, amyotrophic lateral sclerosis; NES, Normalized enrichment score.

To directly compare *Grin2a^+/-^* against *Grin2b*^+/C456Y^, we performed differential expression (DE) and GSEA analysis of 12-week *Grin2a*^+/-^ versus 4-week *Grin2b*^+/C456Y^ bulk RNA-seq, using the associated wild-type mice to correct for batch effects (see Methods). In this comparison, the aforementioned GO terms (such as synapses, OXPHOS, ribosomes) were significantly enriched among the differentially regulated genes (Fig. S1T). Similar results were observed when we performed DE analysis followed up by GSEA in 4-week *Grin2a*^+/-^ versus 4-week *Grin2b*^+/C456Y^ mutant mice (Fig. S1U). Together, these data indicate that there are significant differences in regulation of the same GO terms between *Grin2a*^+/-^ and *Grin2b*^+/C456Y^ mutant mice.

For a more detailed examination of synaptic pathway changes, we conducted GSEA using the refined synaptic ontology terms curated by the SynGO consortium (Koopmans et al., 2019). This analysis uncovered significant changes in both presynaptic and postsynaptic processes, including translation, active zone and postsynaptic density (PSD), in *Grin2a* mutants at 12 weeks, and in *Grin2b*^+/C456Y^ at 4 weeks (Fig. 2B). Strikingly, ‘translation at presynapse’ and ‘translation at postsynapse’ GO terms were upregulated, whereas GO terms related to pre- and postsynaptic structure and organization (presynaptic active zone, PSD) were downregulated in the cortex (PFC/SSC) and hippocampus of *Grin2a* mutants. Of interest, significant downregulation of GO terms related to endocannabinoid signaling, including genes such as *Dagla* and *Plcb1*, was observed in the PFC of *Grin2a*^+/-^ and *Grin2a*^-/-^ mutants (Fig. 2B). This is an intriguing result considering that diacylglycerol lipase alpha (*DAGLA*), a key enzyme involved in endocannabinoid signaling (Schurman et al., 2019), is one of the SCHEMA risk genes with FDR < 5% (Singh et al., 2022). Consistent with this GSEA analysis, multiple genes encoding synaptic proteins were found among the significantly downregulated DEGs in the PFC and hippocampus in *Grin2a*^+/-^ mice, including *Homer1* and *Cacnb4* in the PFC and *Pclo*, *Bsn*, and *Shank2* in the hippocampus (Fig. 2C-D). Like in *Grin2a* mutants, pre- and postsynaptic translation GO terms were significantly upregulated in the cortex and hippocampus of *Grin2b*^+/C456Y^ mice (Fig. 2B). Consistent with this GSEA result, many genes encoding ribosomal proteins (eg *Rps6*, *Rps29*, and *Rpl9*) were among the significantly upregulated DEGs in the hippocampus (Fig. 2E; see Table S2).

We tested whether genes associated with various psychiatric and neurological disorders (see Table S6 for the gene lists) (Buniello et al., 2019; International League Against Epilepsy Consortium on Complex, 2018; Kaplanis et al., 2020; Satterstrom et al., 2020) were enriched among differentially regulated genes in *Grin2a* mutants (Fig. 2F). In both *Grin2a*^+/-^ and *Grin2a*^-/-^, SCZ-associated genes, identified by GWAS and SCHEMA, were significantly enriched among downregulated genes in the PFC, hippocampus, and SSC, and among upregulated genes in the striatum (Fig. 2F). These findings show a link between *Grin2a* mutant mice and the genetics of human SCZ. Additionally, in line with the overlapping genetic basis of SCZ and bipolar disorder (Cross-Disorder Group of the Psychiatric Genomics Consortium, 2019; Palmer et al., 2022), we found a significant enrichment of GWAS hits for bipolar disorder in cortical and hippocampal downregulated genes in *Grin2a* mutants. Unlike SCZ-associated genes, bipolar-associated genes were not enriched among differentially regulated genes of *Grin2a*^+/-^ striatum. Finally, consistent with association of *Grin2a* with neurodevelopmental disorders (NDD) (Endele et al., 2010), and known overlap between SCZ and NDD genes (Singh et al., 2016; Singh et al., 2022), NDD-associated genes were also significantly enriched among downregulated genes in the PFC and hippocampus in *Grin2a* mutants.

### Transcriptomic changes in neuronal and non-neuronal cells in *Grin2a* mutant mice

To identify cell-type specific transcriptomic changes in *Grin2a* mutant mice, we performed single-nucleus RNA-seq (snRNA-seq) in *Grin2a* heterozygous and homozygous mutants and their wild-type littermates at 12 weeks, when we observed the greatest number of DEGs measured by bulk RNA-seq analysis. To allow comparison of cell type-specific changes across ages, we also performed snRNA-seq analysis on PFC and hippocampus from 4-week *Grin2a* animals. No obvious differences in cell type clustering were observed between *Grin2a*^+/-^ and *Grin2a*^-/-^ compared to wild-type mice in any of the tested brain regions at 4 weeks (PFC, hippocampus) or 12 weeks (PFC, hippocampus, SSC, striatum, thalamus) (Fig. S2A, N-R). Observed clusters were annotated to different cell types (Fig. 3A, Fig. S2B-E) based on expression of known marker genes (Fig. S2F-M; Table S7) and comparison to outside datasets.

**Figure 3.**
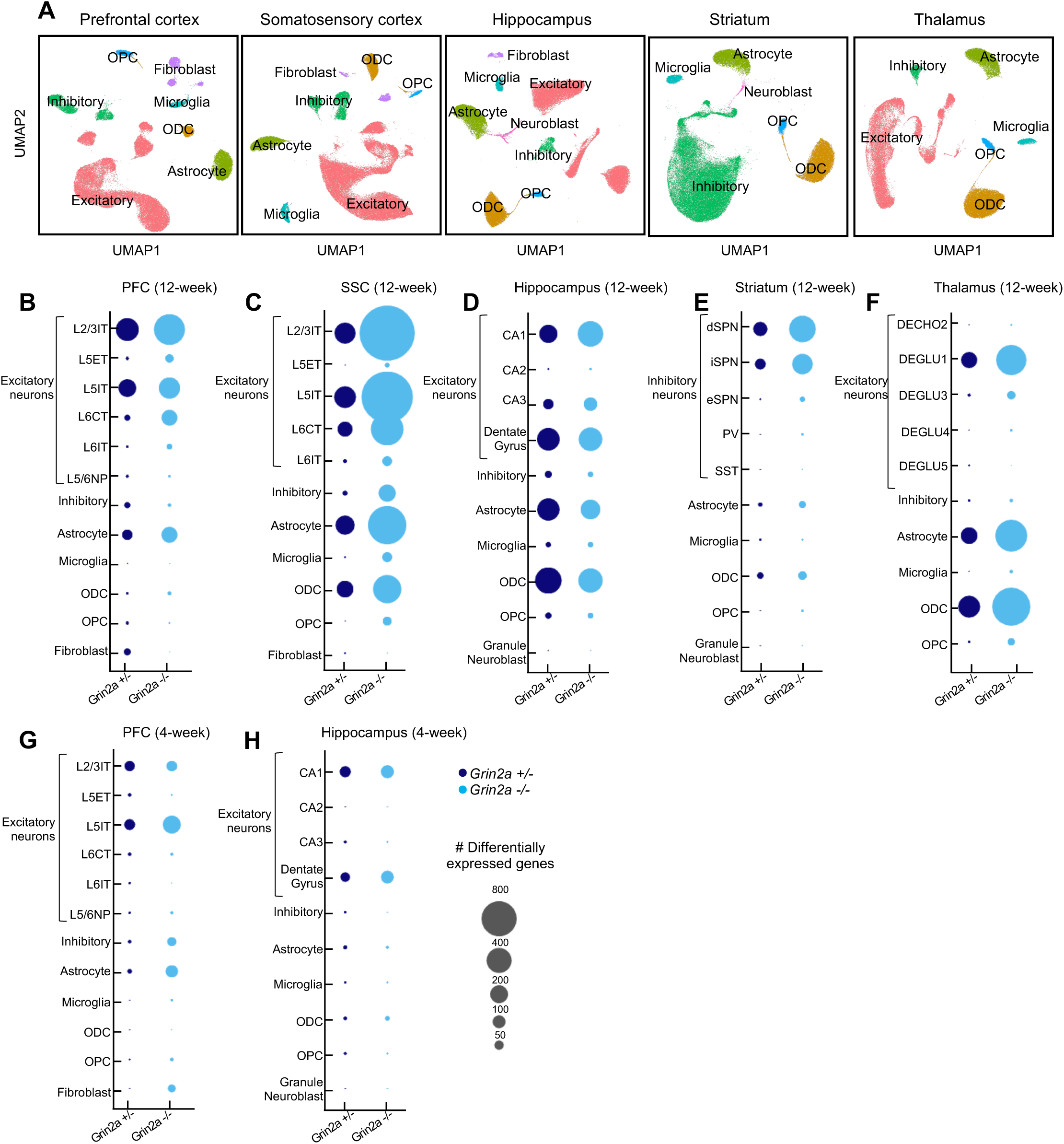
Diverse cell types are affected across brain regions in *Grin2a* mutant mice. (**A**) UMAP representation of the major cell types identified in the five brain regions studied in the *Grin2a* mutant mouse. (**B-H**) Number of DEGs across different cell types in the prefrontal cortex (**B**), somatosensory cortex (**C**), hippocampus (**D**), striatum (**E**) and thalamus (**F**) of 12-week *Grin2a* mutants, and prefrontal cortex (**G**) and hippocampus (**H**) of 4-week *Grin2a* mutants. ODC, oligodendrocyte; PV, parvalbumin interneurons; SST, somatostatin interneurons; VIP, vasoactive intestinal peptide interneurons; L2-6, layers; IT, intratelencephalic; NP, near-projecting; ET, extratelencephalic; CT, corticothalamic neuron; dSPN, direct-pathway spiny projection neurons; iSPN, indirect-pathway spiny projection neurons; eSPN, eccentric spiny projection neurons.

To identify snRNA-seq changes caused by *Grin2a* insufficiency in different cell types, we performed DE analysis using two different algorithms: pseudocell, a mixed model approach that we have recently developed (see Methods), and pseudobulk aggregation followed by EdgeR (Robinson et al., 2010). The quantitative transcriptomic changes (Log2FC values of individual genes) (Fig. S3) as well as the p values were highly correlated between the two approaches (Pearson’s r ≥ 0.8 for Log2FC values and Spearman’s r ≥ 0.6 for p values for most cell types). Also, the consistency scores for both DE analysis and GSEA, defined as the fraction of genes or GO terms that show changes in the same direction in pseudocell and pseudobulk, were greater than 0.9 for most cell types, indicating that the results of the two approaches are largely overlapping albeit detecting different number of DEGs in the same cell types (see Tables S8-14 for pseudobulk DEGs and Tables S22-28 for pseudocell DEGs). Here, we present the results of the pseudocell analysis and provide the DE and GSEA results of pseudobulk analysis for all snRNA-seq datasets in Tables S8-21.

snRNA-seq revealed that multiple cell types, including excitatory and inhibitory neurons as well as non-neuronal cells, were affected in both genotypes of *Grin2a* mutants, across the brain regions and at 4 and 12 weeks of age (Fig. 3B-H). In both *Grin2a*^+/-^ and *Grin2a*^-/-^, excitatory neurons such as cortical (PFC, SSC) pyramidal neurons of layer 2/3 (L2/3IT), and layer 5/6 (L5IT and L6CT; Fig. 3B, C) as well as CA1 pyramidal cells and dentate gyrus excitatory neurons of hippocampus (Fig. 3D) showed large transcriptomic changes at 12 weeks, as measured by number of DEGs. The inhibitory spiny projection neurons (SPNs) of the striatum (Fig. 3E), and DEGLU1 excitatory neurons of thalamus (Fig. 3F) also exhibited large numbers of DEGs at 12 weeks. In 4-week *Grin2a* mutants, there were relatively few DEGs as compared to 12 weeks in different cell types in the PFC and hippocampus (Fig. 3G-H).

Non-neuronal cell types also showed transcriptomic changes by snRNA-seq analysis. Astrocytes and oligodendrocytes (ODCs), in particular, showed a sizable number of DEGs in various brain regions of *Grin2a*^+/-^ and *Grin2a*^-/-^ mice, especially in the SSC, hippocampus and thalamus (Fig. 3B-H). Since glial cells express little or no *Grin2a* mRNA (Fig. S4A), the large transcriptomic changes in astrocytes and ODCs likely reflect indirect (non-cell autonomous) effects of *Grin2a* LoF.

It should be noted that a low count of DEGs in certain cell types might be due to low statistical power of detection in those cell types, rather than lack of transcriptomic changes. For instance, inhibitory interneurons constituted less than 10% of isolated nuclei from cortex (PFC and SSC) and less than 5% of hippocampal nuclei (Fig. S2N-P). The low abundance and high heterogeneity of inhibitory cell types could lead to underestimation of the number of DEGs identified in these cell types by snRNA-seq. In bulk RNA-seq data, we did observe that neuropeptides cortistatin (*Cort)* and neuropepetide Y (*Npy)* and/or somatostatin (*Sst*), which are primarily expressed and released by inhibitory interneurons (Hokfelt et al., 2000; Tremblay et al., 2016), were among the significantly upregulated DEGs (FDR< 0.05) in the hippocampus of *Grin2a*^+/-^ (Fig. 2D) and *Grin2a*^-^

^/-^ (Fig. S4B) and in the PFC and SSC of *Grin2a*^-/-^ (Fig. S4C, D), implying that inhibitory interneurons are also affected by *Grin2a* LoF. Supporting this conclusion, snRNA-seq showed increased expression (positive Log2FC) of a number of neuropeptides including *Npy* and *Cort* in the parvalbumin (PV) and somatostatin (SST) interneurons of the PFC, SSC and hippocampus in *Grin2a*^+/-^ and *Grin2a*^-/-^ (Fig. S4E-P; for a list of neuropeptide genes see Table S6) though these changes did not reach significance at FDR < 0.05.

### Transcriptomic evidence for altered metabolism and PFC hypofunction in *Grin2a* mutant mice

Conducting GSEA in identified cell types in 12-week *Grin2a* mutant mice, we found that OXPHOS-related GO terms (electron transport chain, mitochondrial respiratory complex, etc.) and ribosome-related GO terms showed up commonly as being significantly altered in multiple cell types and brain regions, interestingly not always in the same direction. OXPHOS and ribosomal GO terms were strikingly downregulated in excitatory and inhibitory neurons in the PFC of 12-week *Grin2a*^+/-^ (Fig. 4A, Fig. S5C) and *Grin2a*^-/-^ (Fig. S5A). In contrast, these GO terms were upregulated in neurons of hippocampus, striatum and SSC in *Grin2a* mutants (Fig. 4A, Fig. S5A). Similar region-dependent changes were also observed in 4-week *Grin2a*^+/-^ mice: OXPHOS and ribosomal GO terms were significantly downregulated in diverse subtypes of excitatory and inhibitory neurons in PFC but upregulated in excitatory and inhibitory neuronal subtypes in the hippocampus (Fig. 4B). In *Grin2a*^-/-^ at 4 weeks, however, OXPHOS and ribosomal GO terms were significantly upregulated in excitatory and inhibitory neurons of PFC, which is opposite of what we observed in *Grin2a*^+/-^ mutants (Fig. S5B; see Fig S5D for an example of opposite regulation of ribosomal genes in *Grin2a^+/-^* versus *Grin2a^-/-^* at 4 weeks). These opposite transcriptomic changes related to OXPHOS and ribosomal pathways in *Grin2a*^+/-^ versus *Grin2a*^-/-^ mutants cannot be simply explained by gene dosage effect acting via a cell autonomous mechanism.

**Figure 4.**
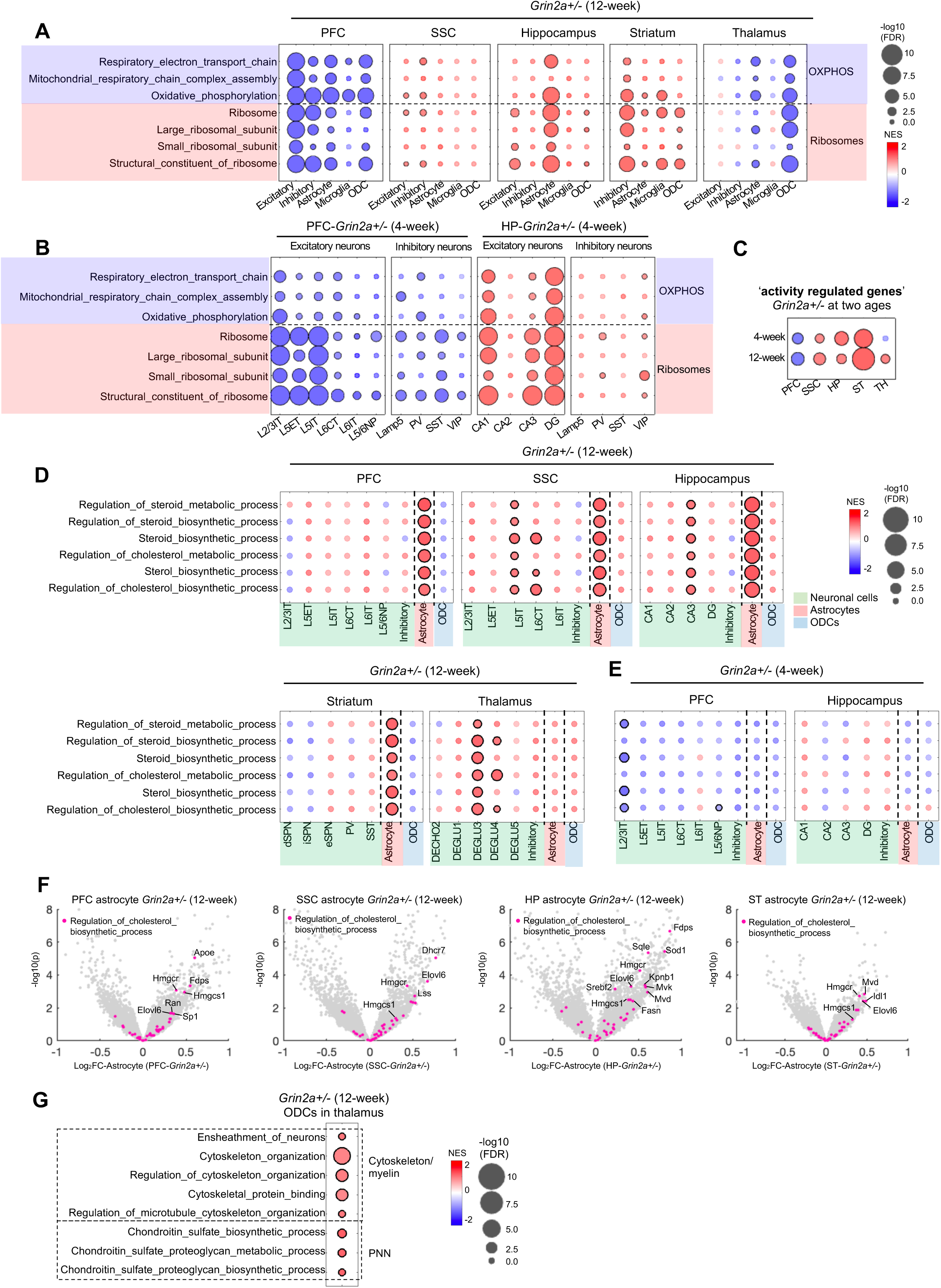
Changes of GO terms in neuronal and non-neuronal cells across brain regions in 4- and 12-week *Grin2a^+/-^* mutant mice. (**A-B**) GSEA results from the 5 major cell types across brain regions in 12-week *Grin2a^+/-^* mutant mice (**A**) and across neuronal subtypes in 4-week *Grin2a^+/-^* mutant mice (**B**). (**C**) Enrichment of the activity-regulated gene set in the whole-tissue transcriptome across brain regions in *Grin2a^+/-^* at 4 and 12 weeks. (**D-E**) GSEA results using GO terms related to cholesterol biosynthesis in excitatory neuronal subtypes, inhibitory neurons, astrocytes and oligodendrocytes (ODCs) across studied brain regions in 12-week (**D**) and 4-week (**E**) *Grin2a^+/-^* mutant mice. (F) Volcano plot of transcriptomic changes in astrocytes of the PFC, SSC, hippocampus and striatum in 12-week *Grin2a^+/-^* highlighting genes from the ‘Regulation of cholesterol biosynthesis process’ GO term. (G) Cytoskeleton/myelin and perineural net (PNN)-related GO terms in thalamic oligodendrocytes in 12-week *Grin2a^+/-^* mutant mice. In all bubble plots circles with black outlines indicate significance (FDR < 0.05). OXPHOS, oxidative phosphorylation.

The changes in OXPHOS (including genes involved in electron transport chain) and ribosomal GO terms suggest a dysregulation of ATP production and protein translation pathways in *Grin2a* mutants which are detectable already at 4 weeks, and differentially affecting PFC, hippocampus and striatum. We hypothesized that the altered expression of OXPHOS- and ribosome-related gene-sets might reflect the metabolic load or metabolic stress on neurons, which should be influenced by the level of activity in those neurons. As a surrogate measure of neuronal activity, we performed GSEA using a curated set of activity-regulated genes (Tyssowski et al., 2018) (see Table S6) in various brain regions. Consistent with our hypothesis, the direction of changes in activity-regulated gene-set agreed well with changes in OXPHOS and ribosomal GO terms in neurons. PFC showed a significant downregulation of the activity-regulated gene-set in *Grin2a*^+/-^ at 4 and 12 weeks (Fig. 4C, Fig. S5E), whereas these genes were significantly enriched among upregulated genes in SSC, hippocampus and especially the striatum (including genes such as *Fos*, *Egr4* and *Chrbp*; Fig. S5F) (Fig. 4C), in line with the directional change of OXPHOS and ribosomal GO terms in these brain regions. Overall, our RNA-seq data support the idea that the widespread, but regionally distinct, changes in neuronal OXPHOS and ribosomal GO terms may be functionally linked to alterations in neuronal activity. Moreover, they spotlight the PFC as a brain region that potentially has reduced activity and decreased ATP production and protein translation in the heterozygous *Grin2a* LoF state, in contrast to other brain regions (SSC, hippocampus, striatum) that show an increase.

In the PFC, non-neuronal cells (astrocytes, microglia and ODCs) also showed downregulation of OXPHOS and ribosomal GO terms in *Grin2a*^+/-^ (Fig. 4A), similar to neurons. Overall, the changes in OXPHOS and ribosome-related GO terms in glial cells were largely correlated with the changes in OXPHOS/ribosomes in neurons of the same brain region (Fig 4A). It has been shown that microglial bioenergetics correlate with their phagocytic activity (He et al., 2022). Downregulation of OXPHOS pathway in microglia, as well as significant upregulation of the “homeostatic” microglia genes *P2ry12*, *Cx3cr1,* and *Olfml3* (Butovsky and Weiner, 2018) in the PFC (see Fig. 2C), suggest that microglia might be in a more homeostatic state with reduced phagocytic activity in the PFC of 12-week *Grin2a*^+/-^ mice.

### Dysregulation of cholesterol biosynthesis in astrocytes and changes in cytoskeletal organization in ODCs of *Grin2a* mutant mice

Unexpectedly, GO terms related to cholesterol/steroid biosynthesis (including genes such as *Hmgcr*, *Hmgcs1*, *Elovl6* and *Srebf2*) were significantly enriched among upregulated genes in astrocytes of the PFC, SSC, hippocampus and striatum in *Grin2a*^+/-^ (Fig. 4D, F) and *Grin2a*^-/-^ mutants (Fig. S5H, J). The upregulation of cholesterol-related GO terms was prominent in astrocytes but was also significant in some subtypes of excitatory neurons in the SSC, hippocampus and PFC (Fig. 4D, Fig. S5H). In thalamus, however, the changes in cholesterol-related GO terms were most prominent in excitatory neurons, notably DEGLU3 excitatory neurons (Fig. 4D, Fig. S5H).

Interestingly, dysregulation of the cholesterol biosynthesis pathway in astrocytes was significant at 12 weeks, but not at 4 weeks (Fig. 4E, Fig. S5I). Consistent with this GSEA analysis, we noted that most cholesterol-related genes showed larger changes at 12 weeks than at 4 weeks in hippocampal astrocytes (Fig. S5K). Thus, the alteration in cholesterol biosynthesis pathway seems to occur relatively late during postnatal brain maturation in *Grin2a* mutant mice.

In ODCs of thalamus, besides OXPHOS and ribosomal GO terms, pathways related to the cytoskeleton and neuron ensheathment (including myelination-related genes such as *Gal3st1*, *Ilk*, *Klk6* (Goudriaan et al., 2014)) were significantly enriched among upregulated genes, suggesting potential myelin abnormalities in the thalamus of *Grin2a*^+/-^ (Fig. 4G). Moreover, GO terms related to the biosynthesis of chondroitin sulfate (which are enriched in the perineural net (PNN)) were significantly upregulated in thalamic ODCs in *Grin2a*^+/-^ (Fig. 4G). Cytoskeleton organization-related pathways were significantly enriched among upregulated genes in *Grin2a*^-/-^ thalamic ODCs as well (Fig. S5L).

### Dysregulation of dopamine signaling in the striatum of *Grin2a* mutant mice

Besides changes in bioenergetics/metabolism pathways (OXPHOS, ribosomes) and neuronal activity (activity regulated genes) that pervaded across multiple brain regions and cell types, there were perturbations in specific neurobiological pathways that affected neuron subtypes and brain regions more selectively. Of particular interest, various GO terms related to dopaminergic and cholinergic signaling were significantly upregulated in the inhibitory neurons of striatum in 12-week *Grin2a*^+/-^ mice (Fig. 5A), ∼95% of which are spiny projection neurons (SPNs; Fig. S2D). SPNs can be categorized as direct spiny projection neurons (dSPN), indirect spiny projection neurons (iSPN), and the recently described striatal eccentric spiny projection neurons (eSPNs) (Saunders et al., 2018). Unlike in *Grin2a*^+/-^ mutants, the GSEA changes in dopamine and cholinergic signaling pathways in striatal neurons did not reach significance in *Grin2a*^-/-^ mice (Fig. S6A).

**Figure 5.**
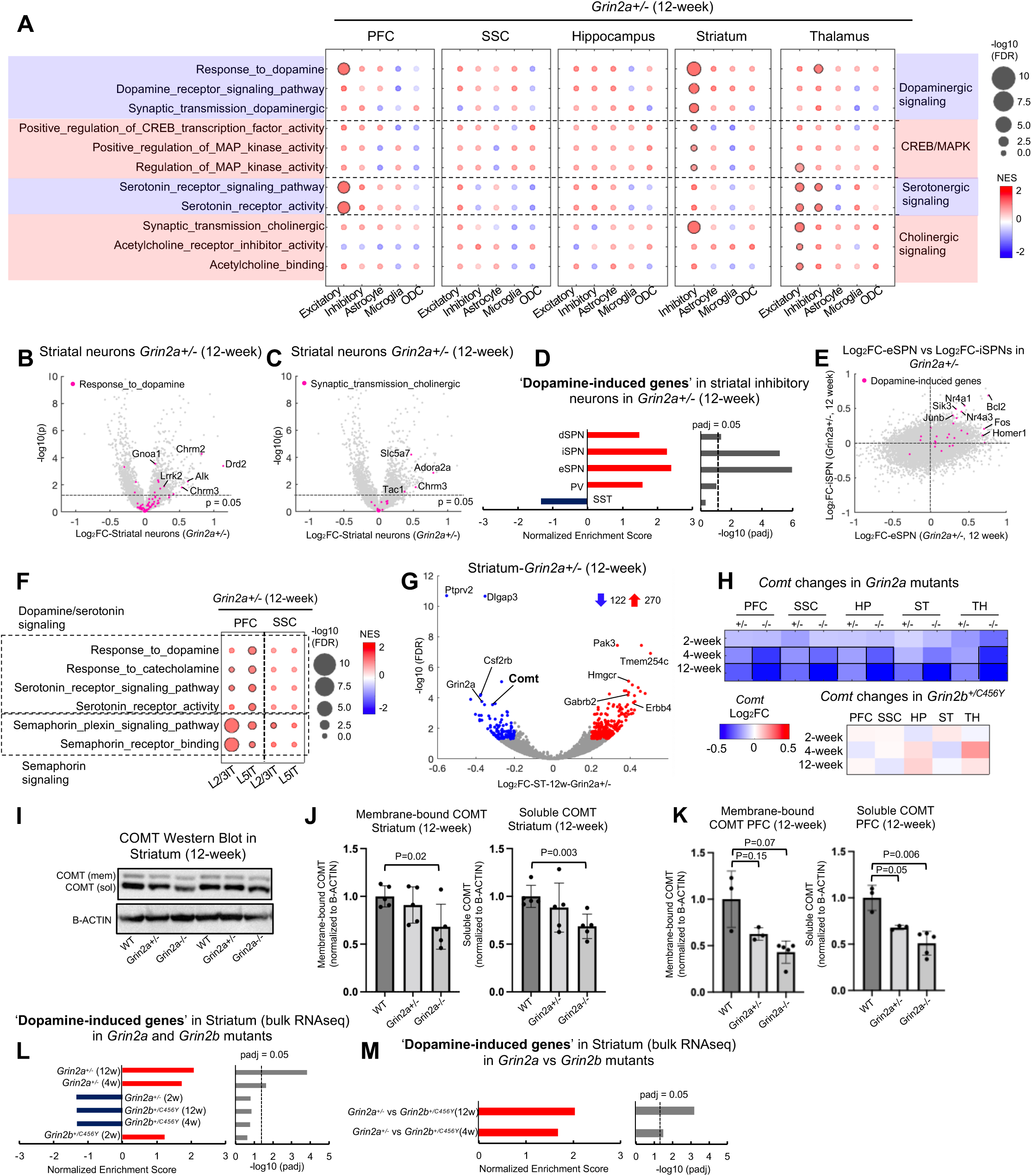
Dysregulation of dopamine in the striatum of *Grin2a^+/-^* mutant mice. (**A**) GSEA results from the 5 major cell types across brain regions of 12-week *Grin2a^+/-^* mutant mice. (**B-C**) Volcano plot of transcriptomic changes in striatal inhibitory neurons in 12-week *Grin2a^+/-^* highlighting genes from ‘Response to dopamine’ (**B**) and ‘Synaptic transmission cholinergic’ (**C**) (D) Enrichment of the dopamine-induced gene-set in differentially regulated genes of striatal inhibitory neuron subtypes in 12-week *Grin2a^+/-^*. (E) Transcriptomic changes in eSPNs versus iSPNs in the striatum of 12-week *Grin2a^+/-^* highlighting the dopamine-induced gene set. (F) Changes in serotonin, dopamine and semaphorin signaling GO terms in layer 2/3IT and layer 5IT excitatory neurons in the PFC and SSC of 12-week *Grin2a^+/-^*. (G) Volcano plots showing all significant DEGs (blue and red circles represent down- and upregulated DEGs, respectively) in the striatum of 12-week *Grin2a^+/-^*. Numbers next to blue and red arrows indicate the number of significantly down- and upregulated DEGs, respectively. Genes of interest are labeled on the volcano plot. (H) Heatmap showing the Log2FC values of *Comt* in the bulk RNA-seq data from studied brain regions in 2-, 4- and 12-week *Grin2a* and *Grin2b* mutant mice. Black outlines indicate significance (FDR < 0.05). (I) Western blots probing for COMT and B-ACTIN in total striatal lysate obtained from 12-week *Grin2a* animals. (**J-K**) Quantification of COMT protein expression measured by Western blot in the striatum (**J**) and PFC (**K**) of *Grin2a* animals. Data are shown as mean +/- standard error (n=3-5 animals per genotype). p values are computed using two-tailed Student’s t test. (L) Enrichment of the dopamine-induced gene-set in differentially regulated genes of the striatum (bulk RNA-seq) in *Grin2a^+/-^* and *Grin2b^C456Y/+^* at 2, 4 and 12 weeks. (M) Enrichment of the dopamine-induced gene-set in differentially regulated genes of the striatum (bulk RNA-seq) in *Grin2a^+/-^* versus *Grin2b^C456Y/+^* DE analysis at 4 and 12 weeks. In all bubble plots circles with black outlines indicate significance (FDR < 0.05). CREB, cyclic adenosine 3′,5′-monophosphate (cAMP) response element–binding protein; MAPK, mitogen-activated protein kinase; WT, wild type.

Multiple genes within the ‘Response_to_dopamine’ and ‘Synaptic_transmission_cholinergic’ GO terms (e.g. *Chrm3*, *Slc5a7*, *Chrm2*, *Drd2* and *Lrrk2*) were elevated in striatal inhibitory neurons of 12-week *Grin2a*^+/-^ (Fig. 5B, C). It did not escape our attention that *Drd2* (D2 dopamine receptor, the target of most anti-psychotic drugs), and *Chrm2* (M2 muscarinic receptor), were among the most highly and significantly upregulated DEGs in the striatal inhibitory neurons of 12-week *Grin2a*^+/-^ mice (Fig. 5B).

The transcription factor CREB (cAMP response element–binding protein) and its activation via MAPK (mitogen-activated protein kinase) pathways are believed to play a vital role in dopamine receptor signaling (Andersson et al., 2001). Consistent with enhanced dopamine signaling, GO terms related to positive regulation of CREB activity and MAPK pathways were significantly upregulated in the striatal neurons of *Grin2a*^+/-^ mice (Fig. 5A). Moreover, dSPN and iSPNs in *Grin2a*^+/-^ showed an increased expression of several cAMP-induced CREB target genes (Zhang et al., 2005), including *Bcl2* (Fig. S6B; Table S6), whose expression rises in rat SPNs in response to increased dopamine release elicited by cocaine treatment (Savell et al., 2020). Extending these results, we found a significant enrichment of a set of dopamine-induced genes (Table S6; including activity-regulated genes *Homer1*, *Fos* and *Junb* that are induced by dopamine (Savell et al., 2020)) among the upregulated genes of *Grin2a*^+/-^ striatum (Fig. 5D). Notably, the upregulation of the dopamine-induced gene-set was strong (NES > 2) and significant in iSPNs which express mainly Drd2 dopamine receptors, and in eSPNs in which we observed expression of both Drd2 and Drd1 dopamine receptors (see Methods) (Fig. 5D, E).

We also found evidence for dopamine and serotonin signaling changes in the PFC of 12-week *Grin2a*^+/-^ mice, where dopamine/serotonin signaling GO terms were significantly enriched among upregulated genes in L2/3IT and L5IT pyramidal cells (Fig. 5F). These same cell types also showed significant upregulation of GO terms related to semaphorin-plexin signaling (Fig. 5F), which is known to be involved in the remodeling of synaptic connections (O’Connor et al., 2009; Simonetti et al., 2021; Uesaka et al., 2014) and has been implicated in SCZ (Eastwood et al., 2003; Zhou et al., 2012). In line with this GSEA result, plexins *Plxnb1* and *Plxnb2* as well as semaphorin *Sema3g* were found among the significantly upregulated DEGs in the PFC of 12-week *Grin2a*^+/-^ mice (see Fig. 2C). Changes in most of these GO terms related to dopamine/serotonin/semaphorin signaling did not reach significance in the SSC of *Grin2a*^+/-^ (Fig. 5F), or in PFC and SSC of *Grin2a*^-/-^ mice (Fig. S6C).

In the context of heightened dopamine signaling, it is noteworthy that *Comt* (encoding catechol-O-methyltransferase, an enzyme involved in dopamine degradation) was among the most significantly downregulated genes in the bulk RNA-seq analysis of striatum in 12-week *Grin2a*^+/-^ (Fig. 5G). In fact, *Comt* mRNA was consistently reduced in a gene dose-dependent fashion in all tested brain regions of *Grin2a* mutants (Fig. 5H). In sharp contrast, *Grin2b^+/C456Y^* mutants showed no significant changes in *Comt* mRNA in any of the studied brain regions (Fig. 5H). We confirmed reduction of COMT protein by Western blotting; compared with wild types, COMT levels fell by ∼50% in the PFC and ∼40% in the striatum in 12-week *Grin2a*^-/-^, with *Grin2a*^+/-^ showing intermediate reduction (Fig. 5I-K). Decreased expression of *Comt* might lead to elevated dopamine levels and contribute to the enhanced expression of dopamine-regulated genes in the striatum of *Grin2a^+/-^* (Fig. 5D). Consistent with such a notion, we observed a significant enrichment of the dopamine-induced gene-set among upregulated genes of striatum in *Grin2a*^+/-^ mutants (Fig. 5L). This was seen at 4 and 12 weeks, but not at 2 weeks, suggesting that striatal dopamine dysregulation emerges during the juvenile period of brain maturation in *Grin2a^+/-^* mice (Fig. 5L). In *Grin2b^+/C456Y^* mutants, however, no significant enrichment of the dopamine-induced gene-set was observed among differentially regulated genes at any of the tested ages (Fig. 5L), in line with the unchanged *Comt* expression in *Grin2b^+/C456Y^* mutants (Fig. 5H). In a direct DE analysis comparison between the two genotypes, the dopamine-induced gene-set was significantly upregulated in *Grin2a^+/-^* versus *Grin2b^+/C456Y^* mutants (Fig. 5M). Together, these data point to dysregulated dopamine signaling, suggestive of a hyperdopaminergic state, in the striatum and PFC of specifically *Grin2a* mutant mice.

In the thalamus of *Grin2a^+/-^*, serotonin and cholinergic signaling GO terms were found significantly upregulated in the excitatory neurons of *Grin2a*^+/-^ (Fig. 5A). A significant upregulation of some dopaminergic/serotoninergic signaling pathways was also observed in the inhibitory neurons of the thalamus in both *Grin2a*^+/-^ (Fig. 5A) and *Grin2a*^-/-^ (Fig. S6A). Of interest, *Htr2a* and *Htr2c* (encoding serotonin receptors 2A and 2C) were among the most upregulated genes in the ‘Serotonin_receptor_signaling_pathway’ GO term in the thalamic inhibitory neurons of *Grin2a*^+/-^ (Fig. S6D).

### Changes in glutamatergic signaling and ribosomal proteins in synaptic proteome of *Grin2a* mutant mice

Our RNA-seq GSEA analysis point to major changes in synapse-related pathways in *Grin2a* mutant brains (Fig. 2A, B). To investigate the effects of *Grin2a* LoF on synapses at the protein level, we conducted quantitative mass spectrometry (MS) proteomics of synaptic fractions, purified from the hippocampus of 4-week *Grin2a*^+/-^, *Grin2a*^-/-^ and their wild-type littermates (see Methods). We observed a similar number of differentially expressed proteins (DEPs, defined as proteins with significant changes (p < 0.05 and absolute values of Log2FC > 0.2) versus wildtype) in *Grin2a*^+/-^ (Fig. 6A; 291 DEPs) and *Grin2a*^-/-^ (Fig. 6B; 282 DEPs). The vast majority (97%) of the significant DEPs in *Grin2a*^+/-^ were altered in the same direction in *Grin2a*^-/-^ (Fig. 6C), such as XPO7, GRIN3A and RPL37 (Fig. 6A, B). Evaluation of the Log2FC values of all individual proteins in *Grin2a*^+/-^ versus *Grin2a*^-/-^ synapses revealed an overall strong correlation and a similar magnitude of change of the synaptic proteomes in *Grin2a*^+/-^ and *Grin2a*^-/-^ (Pearson’s r = 0.75; Fig. 6D). Overall, these data signify that heterozygous *Grin2a* LoF has a similar-size effect on global synapse composition as homozygous LoF, which is in line with our conclusion from bulk RNA-seq data that *Grin2a*^+/-^ has comparable effect as *Grin2a*^-/-^ on the transcriptome (Fig. 1B)

**Figure 6.**
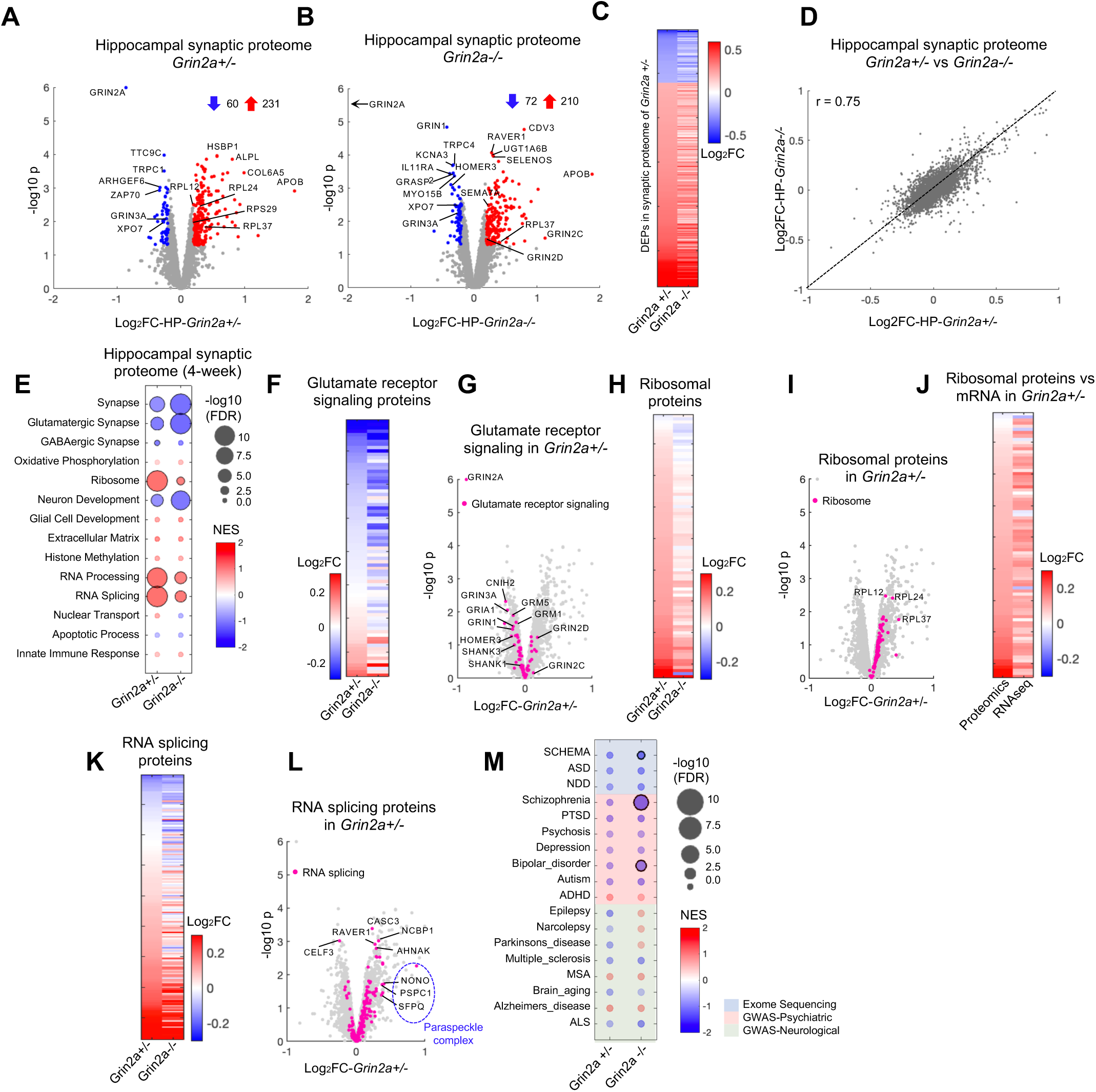
Synapse proteome changes in *Grin2a* mutant mice. (**A-B**) Volcano plots showing significant DEPs (defined as proteins with changes versus wild types with nominal p < 0.05 and absolute value of Log2FC > 0.2; blue and red circles represent down- and upregulated DEPs, respectively) in the hippocampal synapses of 4-week *Grin2a^+/-^* (**A**) and *Grin2a^-/-^* (**B**). Numbers next to blue and red arrows indicate the number of significantly down- and upregulated DEPs, respectively. (C) Heatmap showing Log2FC values in both *Grin2a^+/-^* and *Grin2a^-/-^* for significant DEPs identified in the hippocampus of *Grin2a^+/-^*. (D) Correlation of protein changes in the hippocampus of *Grin2a^+/-^* versus *Grin2a^-/-^*. Pearson’s r correlation value is indicated on the plot. (E) GSEA results in the hippocampal synapse proteome of 4-week *Grin2a^+/-^* and *Grin2a^-/-^*. (F) Heatmap depicting all detected proteins in the ‘glutamate receptor signaling pathway’ gene-set in the hippocampal synaptic proteome of 4-week *Grin2a^+/-^* and *Grin2a^-/-^*. (G) Volcano plot of proteomic changes in *Grin2a^+/-^* hippocampal synapses highlighting proteins in the ‘glutamate receptor signaling pathway’ gene-set. (H) Heatmap depicting all detected proteins in the ‘cytosolic ribosome’ gene-set in the hippocampal synaptic proteome of 4-week *Grin2a^+/-^* and *Grin2a^-/-^*. (I) Volcano plot of proteomic changes in *Grin2a^+/-^* synapses highlighting proteins in the ‘cytosolic ribosome’ gene-set. (J) Heatmap depicting all detected proteins in the ‘cytosolic ribosome’ gene-set in the hippocampal synaptic proteome and their mRNA changes in hippocampal bulk RNA-seq data in 4-week *Grin2a^+/-^*. (K) Heatmap depicting all detected proteins in the ‘RNA splicing’ gene-set in the hippocampal synaptic proteome of 4-week *Grin2a^+/-^* and *Grin2a^-/-^*. (L) Volcano plot of proteomic changes in *Grin2a^+/-^* synapses highlighting proteins in the ‘RNA splicing’ gene-set. (M) GSEA results in the hippocampus of 4-week *Grin2a^+/-^* and *Grin2a^-/-^* showing enrichment patterns for risk variants from psychiatric and neurological diseases identified by exome sequencing and GWAS studies (abbreviations as in Fig. 2). In **B** arrow indicates that GRIN2A is out of the range of the plots. In **E** and **M**, circles with black outlines indicate significance (FDR < 0.05). In **C**, **F**, **H** and **K**, each row represents a protein, and the column represents the genotype of the animal. Rows are sorted in ascending order based on the Log2FC values from *Grin2a^+/-^*.

To validate the MS results, we measured several synaptic proteins (including GRIN2A itself) by Western blotting of hippocampal synaptic fractions purified from an independent mouse cohort (Fig. S7A). The magnitude and direction of the effect sizes we observed by Western were highly consistent with those measured by MS (Fig. S7B).

GSEA of the proteomics data showed that GO terms related to synapses, particularly glutamatergic synapses, were significantly downregulated in synapses of *Grin2a* mutants (Fig. 6E). Perhaps not surprising since GRIN2A is a protein of glutamatergic synapses, the magnitude of reduction of many downregulated proteins of glutamatergic signaling was greater in *Grin2a*^-/-^ than in *Grin2a*^+/-^ (Fig. 6F). Notably, many glutamate receptor subunits (e.g. GRIN1, GRIN3A, GRM5, GRIA1 and GRIA2), and the scaffolding proteins SHANK1, SHANK3, and HOMER3 were among the downregulated proteins in both *Grin2a*^+/-^ (Fig. 6G) and *Grin2a*^-/-^ (Fig. S7C). We did notice an increase in NMDA receptor subunit GRIN2D, which might be a compensatory response to GRIN2A deficiency (Fig. 6G, Fig.S7C).

More unexpected than downregulation of glutamatergic signaling proteins was the finding that GO terms related to ribosomes and RNA processing/splicing were significantly upregulated in *Grin2a* mutants (interestingly, to a greater degree in *Grin2a*^+/-^ synapses than in *Grin2a*^-/-^) (Fig. 6H, K). Ribosomal proteins showed striking elevation as a group in the synapses of *Grin2a*^+/-^ (Fig. 6I) and, to a lesser extent, *Grin2a*^-/-^ mutant mice (Fig. S7D). The increase in ribosomal proteins in the synaptic proteome is consistent with the upregulation of ribosomal gene expression in bulk RNA-seq data from 4-week hippocampus (Fig. S7E, F) and in snRNA-seq data from excitatory neurons and inhibitory interneurons in the hippocampus of 4-week *Grin2a* mutants (Fig. 4B, Fig. S5B). In fact, more than 90% of ribosomal proteins that were upregulated in the synaptic proteome of *Grin2a*^+/-^ and *Grin2a*^-/-^ were also upregulated at the mRNA level in bulk RNA-seq analysis of the hippocampus in 4-week *Grin2a* mutant brains (Fig. 6J, Fig. S7G). However, the correlation of the Log2FC values of mRNA and synaptic protein level of these ribosomal genes was only modest (r = 0.2) in *Grin2a*^+/-^ and *Grin2a*^-/-^ hippocampus, suggesting that post-transcriptional changes, such as protein turnover and trafficking, likely contribute to ribosomal protein changes at synapses in *Grin2a* mutant mice. The upregulation of ribosomes in the synaptic proteome and the increased mRNA of ribosomal genes point to potentially increased protein synthesis at synapses, an interpretation that is corroborated by SynGO analysis (Fig. 2B). Enhanced translation has been observed in neurons with mutations in *Fmr1* gene (Aryal et al., 2021), and excessive protein synthesis has been causally linked to the phenotypes of Fragile X Syndrome (Richter et al., 2015). Surprisingly, we found that more than 60% of the identified RNA splicing proteins were elevated in *Grin2a*^+/-^ (Fig. 6L) and *Grin2a*^-/-^ synapses (Fig. S7H). Among these, proteins of the paraspeckle complex (SFPQ, NONO, and PSPC1) were prominently upregulated in both *Grin2a*^+/-^ and *Grin2a*^-^

^/-^ synapses. Paraspeckles are subnuclear bodies found in the interchromatin space of mammalian cell nuclei and are involved in several aspects of RNA processing, including transcription initiation, transcriptional termination, and mRNA splicing (Fox and Lamond, 2010). The presence and increased abundance of the paraspeckle complex and other RNA processing/splicing proteins in synaptic fractions of *Grin2a* mutants might indicate mislocalization of these RNA binding proteins from the nucleus to the cytoplasm, a pathomechanism that has been implicated in neurodegenerative diseases such as ALS and frontotemporal dementia (Lester et al., 2021; Lim et al., 2020).

We also conducted GSEA on the synapse proteome for gene sets that are associated with various psychiatric and neurological disorders (Table S6): SCZ-associated genes identified by GWAS and SCHEMA were significantly enriched among downregulated proteins at *Grin2a*^-/-^ synapses (Fig. 6M). Furthermore, similar to our results from bulk RNA-seq analysis (Fig. 2F), bipolar disorder GWAS genes were enriched among the downregulated proteins of *Grin2a*^-/-^ hippocampal synapses.

### Abnormal EEG and locomotor behavior of *Grin2a* mutant mice

To investigate how *Grin2a* LoF affects brain network activity, we monitored EEG in *Grin2a* mutant mice, analyzing signals from an electrode over the parietal cortex (Fig. 7A-E). Full details of EEG recordings in *Grin2a* mutant mice including stimulus-evoked EEG measurements, will be described elsewhere (see (Herzog et al., 2022)). At ∼3 months of age, *Grin2a*^+/-^ and *Grin2a*^-/-^ mutants showed a significant increase in gamma (30-50 Hz) oscillation power during non-rapid eye movement (NREM) sleep (by ∼10% and ∼15%, respectively; Fig. 7A). This elevation in resting gamma power in *Grin2a* mutant mice is reminiscent of the increase in gamma oscillations during sleep and quiet wake in SCZ patients (Tanaka-Koshiyama et al., 2020; Tekell et al., 2005). *Grin2a*^-/-^ also displayed elevated slow, delta, sigma and beta oscillations relative to their wild-type littermates (∼10-15% increase), while *Grin2a*^+/-^ was not significantly different from wild type at these frequency bands (Fig. 7B-E). Theta and alpha oscillations were similar across all genotypes (Fig. S8A, B).

**Figure 7.**
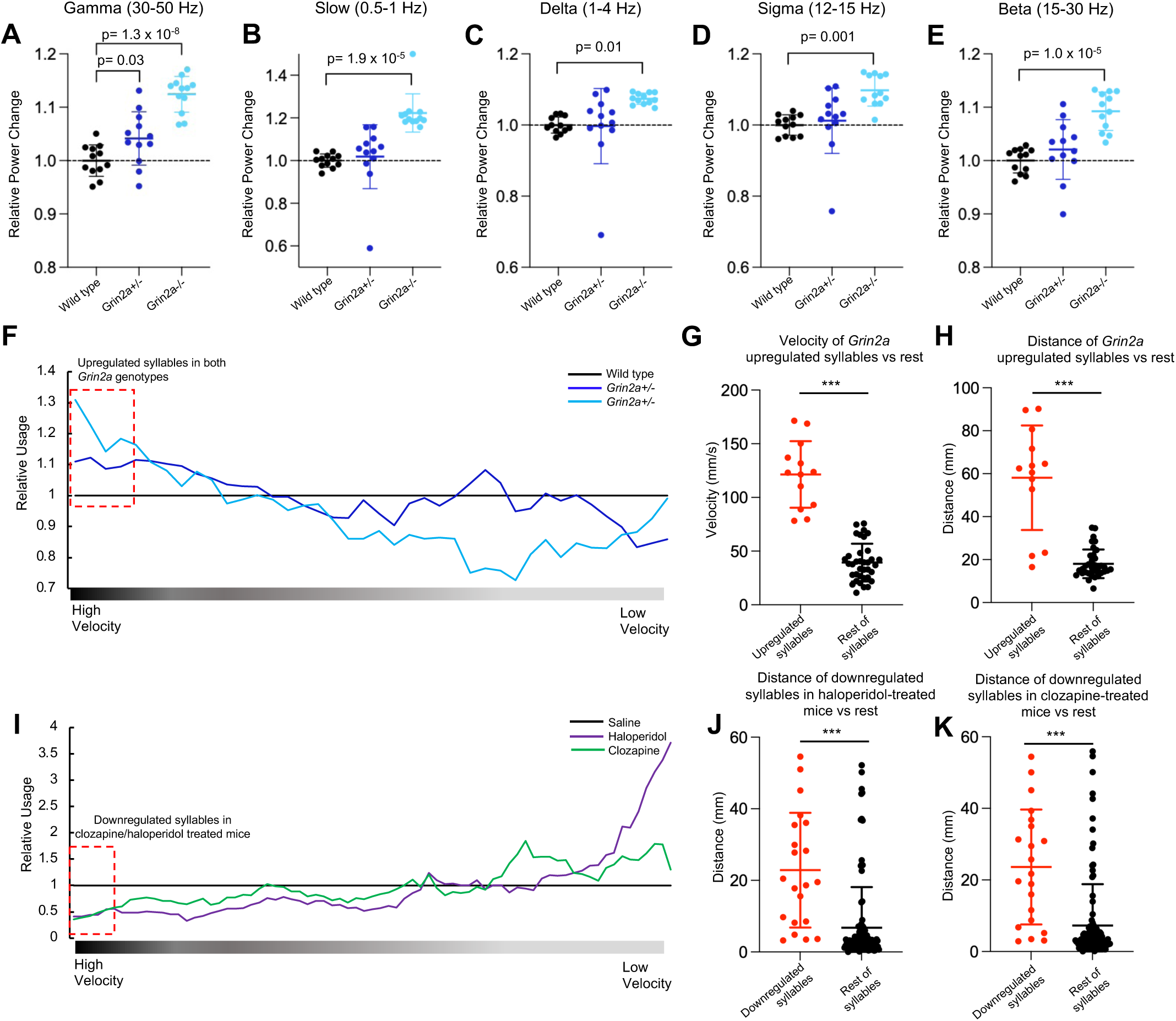
Abnormal EEG and locomotor behavior of *Grin2a* mutant mice. (**A-E**) Power spectral density (PSD) changes in *Grin2a* mutant mice relative to wild types at ∼3 months of age, measured during NREM sleep during the light cycle for each of the frequency bands: (**A**) gamma, 30-50 Hz; (**B**) slow, 0.5-1 Hz; (**C**) delta, 1-4 Hz; (**D**) sigma, 12-15 Hz; (**E**) beta, 15-30 Hz. Error bars show mean +/- standard error; n=12 mice/group; p values were computed using one-way ANOVAs with post hoc Tukey-Kramer tests for multiple comparisons. WT, wild type. (**F**) Relative average syllable usage in *Grin2a^+/-^* and *Grin2a^-/-^* normalized to wild types, after sorting the syllables by mean velocity from high to low and averaging them with a sliding window of 10 syllables. Red box indicates top quartile syllables with highest velocity which are upregulated in both *Grin2a* genotypes. (**G-H**) Mean velocity within syllables (**G**) and mean distance traveled during syllables (**H**) for top quartile syllables which showed upregulation in both *Grin2a* mutants (red box in **F**) versus rest of identified syllables. (**I**) Relative average syllable usage in wild-type mice treated with either clozapine (10mg/kg) or haloperidol (0.25 mg/kg) normalized to saline-treated mice after sorting the syllables by mean velocity from high to low and averaging them with a sliding window of 10 syllables. Red box indicates top quartile syllables with highest velocity which are downregulated in clozapine/haloperidol treated animals. (**J-K**) Mean distance traveled during downregulated syllables in haloperidol- (**J**) and clozapine- (**K**) treated mice (red box in **I**) versus rest of identified syllables. For **I-K** data are taken from (Wiltschko et al., 2020). In **G**, **H**, **J** and **K** error bars show mean +/- standard error. Asterisks (*) indicate statistical significance assessed using non-parametric permutation tests (see Methods); (***) indicates p < 0.001.

We measured behavior by motion sequencing (MoSeq) analysis, an unsupervised machine learning framework that characterizes mouse behavior in the format of subsecond behavioral motifs, referred to as syllables (Wiltschko et al., 2020). Both kinematic values (see Methods) and MoSeq behavioral summaries were computed in the pipeline for each animal (Fig. S8C, D). No significant differences were observed between *Grin2a*^+/-^ mutants and their wild-type littermates in kinematic measurements such as size of the animals, their mean velocity, total distances traveled, and their mean distance to the arena center (Fig. S8E-H). *Grin2a*^-/-^ mutants showed higher mean velocity than wild type (Fig. S8F), but no other kinematic values were significantly different.

By sorting the behavioral syllables by their velocity and comparing usage in *Grin2a* mutants versus wild-type animals, however, we noted that the top quartile with highest-velocity syllables were used more by the *Grin2a* mutant mice as compared to wild types (Fig. 7F). The syllables of the top quartile (shown with a red box in Fig. 7F) showed significantly greater velocity that the rest of identified syllables (Fig. 7G). These same syllables also showed significantly greater mean distances traveled during the syllables when compared to the rest of the syllable population (Fig. 7H). Collectively, these data suggest that *Grin2a* mutant mice perform movements with high locomotor activity and velocity more often than their wild-type littermates. These behavioral patterns identified with MoSeq extend the previously reported increased locomotor activity of *Grin2a* mutant mice (Boyce-Rustay and Holmes, 2006; Hanson et al., 2020; Herzog et al., 2022). Locomotor hyperactivity in open field has been reported in other animal models of glutamate hypofunction (including pharmacologically-induced models), and can be ameliorated by treatment with antipsychotic drugs (Corbett et al., 1995; Moghaddam and Adams, 1998; Mohn et al., 1999). To further contextualize our MoSeq findings, we analyzed previously published datasets collected from wild-type mice treated with two antipsychotic drugs, haloperidol (0.25 mg/kg) or clozapine (10 mg/kg) (Wiltschko et al., 2020), and evaluated how syllable usage is affected after these treatments. Interestingly, the syllables with highest velocity were downregulated in mice treated with haloperidol or clozapine (Fig. 7I). Additionally, the top quartile syllables with highest velocity (shown with a red box in Fig. 7I) showed significantly greater velocity (Fig. S8I, J), and significantly greater mean distances traveled during the syllables when compared to the rest of the syllable population (Fig. 7J, K). These data show that treatment of mice with haloperidol or clozapine results in less usage of movements which involve higher locomotor activity and greater velocity, a behavioral pattern that is opposite to what we observed in *Grin2a* mutant mice.

## DISCUSSION

An animal model that has human-genetics validity, and that exhibits key neurobiological features of the human disease, would have transformative impact on mechanistic understanding of SCZ. Heterozygous LoF mutations that strongly elevate the SCZ risk in humans, such as those in *GRIN2A* (Singh et al., 2022), offer the opportunity to create such models through heterozygous disruption of the gene in mice and other animals. Here, studying the heterozygous *Grin2a* mutant mouse at multiple levels, and with multi-omics approaches, we made a series of discoveries that open new windows into schizophrenia mechanism.

First, even though *Grin2a* and *Grin2b* encode subunits of the same NMDARs, the transcriptomic changes in *Grin2a^+/-^* brain were poorly or even anti-correlated with the same brain region in *Grin2b^+/C456Y^* mice (Fig. 1F-H, Fig. S1O, P). The distinct global transcriptomic changes are consistent with *Grin2a* and *Grin2b* having non-identical spatiotemporal expression patterns in the brain (Paoletti et al., 2013) and different disease associations: *GRIN2A* but not *GRIN2B* is associated with schizophrenia (Platzer et al., 2017; Singh et al., 2022). We also noted that in general the RNA-seq changes in the *Grin2b^+/C456Y^* mutant brain are bigger at 4 weeks than at 12 weeks of age while in most brain regions in *Grin2a^+/-^* mutants, especially in the striatum, bigger transcriptomic changes occurred at 12 weeks of age (Fig. S1F), a period of brain maturation which might be relevant to the typical adolescence/young adult onset of SCZ.

Second, *Grin2a^+/-^* mice show surprisingly large changes in transcriptomic and proteomic phenotype, comparable to, and sometimes even greater than, *Grin2a^-/-^* mutants. For instance, the overall RNA-seq profile of PFC changed to a greater degree in *Grin2a^+/-^* than in *Grin2a^-/-^* (Fig. 1C), and ribosomal proteins were more elevated in the hippocampal synaptic proteome of *Grin2a^+/-^* than in *Grin2a^-/-^* (Fig. 6H). In line with these results, loss of just one copy of *Grin2a* is sufficient to cause behavioral change (hyperactivity in MoSeq) and neurophysiological abnormality (elevated gamma oscillation power on EEG), both of which are features of human SCZ and/or pharmacological models of SCZ (Tanaka-Koshiyama et al., 2020; Tekell et al., 2005; van den Buuse, 2010), see also (Herzog et al., 2022).

Third, *Grin2a* LoF has markedly different effects on different brain regions, exemplified by significant changes in synapse-related GO terms in *opposite* directions in striatum versus cortex (PFC/SSC) and hippocampus (Fig. 2A, Fig. S1Q). It is also remarkable that measures of neuronal activity (activity regulated gene-set) and neuronal metabolism (OXPHOS/mitochondrial respiration, ribosomes) which were correlated with each other within brain regions, moved in opposite directions in different brain regions. Activity-, OXPHOS- and ribosome-related gene-sets were consistently and robustly downregulated specifically in the PFC in *Grin2a^+/-^* mice, whereas these same activity and metabolism GO terms were upregulated in the striatum and hippocampus (Fig 4A, C). The PFC, a brain region involved in higher cognitive and executive functions, has been much studied in relation to cognitive impairment in SCZ (Sakurai et al., 2015; Sigurdsson et al., 2010). In this context, it is interesting that *Grin2a^+/-^* impacts the PFC much more than heterozygous LoF of the ASD gene *Grin2b* (Fig. 1F), which is consistent with a recent human brain transcriptomic study showing that ASD has relatively little effect on PFC compared to other neocortical areas (Gandal et al., 2022). Our finding, based on transcriptomics, that *Grin2a^+/-^* mutants exhibit hypofunction particularly in PFC is intriguing because there is clinical evidence for reduced activity and brain volume in parts of PFC in SCZ by functional neuroimaging (Minzenberg et al., 2009; Pomarol-Clotet et al., 2010). Further correlating with our RNA-seq observations (Fig. 4A-C), SCZ patients show hyperactivity in hippocampus (Holt et al., 2005; Malaspina et al., 1999; Schobel et al., 2013; Talati et al., 2014; Tregellas et al., 2009; Tregellas et al., 2014), which correlates with symptoms of psychosis (Silbersweig et al., 1995).

Fourth, among many pathway changes observed, perhaps most impactful is that *Grin2a^+/-^* mice show evidence of hyperdopaminergic signaling in the striatum and PFC. GO terms related to dopamine response and signaling were significantly upregulated in pyramidal neurons of PFC and spiny projection neurons of striatum, while levels of *Comt* mRNA and protein fell (Fig. 5). Increased dopamine release has been observed in the PFC and striatum following treatment with NMDAR antagonists (Loscher et al., 1991; Moghaddam et al., 1997). In clinical studies, a hyperdopaminergic state of the striatum has been reported in SCZ patients (Grace, 2016; Laruelle and Abi-Dargham, 1999; Laruelle et al., 1999), possibly driven by increased activity in hippocampal efferents that is postulated to elevate dopaminergic input into striatum (Stone et al., 2010). Hyperdopaminergic signaling, particularly in the striatum, has been a long-standing hypothesis for the pathophysiology of schizophrenia, not least because anti-psychotic drugs (which show considerable efficacy against the positive symptoms of schizophrenia) have antagonist activity against the D2 dopamine receptor (Kapur and Mamo, 2003; Meltzer et al., 2003). In this light, it is striking that in *Grin2a^+/^*^-^ mice, one of the most highly elevated transcripts in striatal SPNs was *Drd2* (Log2FC = 1.13; FDR = 0.03), which encodes the D2 dopamine receptor (Fig. 5B). Striatal dopamine abnormalities have been postulated to be secondary to circuit dysfunction caused by altered glutamate signaling in inhibitory neurons during early postnatal brain development (Nakazawa and Sapkota, 2020). Consistent with the hyperdopamingergic state being a later secondary response, changes in dopamine-induced gene-set emerged at 4 weeks of age in the striatum of *Grin2a^+/-^* mutants (Fig. 5L), whereas changes in GO terms related to synapses and neuronal metabolism (OXPHOS, ribosomes) were already apparent at 2 weeks of age (see Table S4). Moreover, our transcriptomic data provide strong evidence that the functional state (though not the number) of inhibitory interneurons is affected by *Grin2a* heterozygous LoF, including alteration of OXPHOS and ribosomal GO terms in inhibitory interneurons (Fig. 4A, B), and elevation of neuropeptides expression such as *Cort* and *Npy* in interneurons of cortex and hippocampus (Fig. S4E-P). Together, our data suggest that dopamine dysregulation in *Grin2a^+/-^* mutants may be a downstream event of NMDAR hypofunction in the brain, perhaps involving both excitatory and inhibitory neuron dysfunction. Although the mechanistic link between *Grin2a* LoF and hyperdopaminergic drive in the striatum is not clear, it is impressive that heterozygous mutation in *Grin2a*, a gene that surfaced from human genetics of schizophrenia, marries the two major prevailing hypotheses of schizophrenia pathophysiology: (i) the “NMDAR hypofunction” hypothesis, and (ii) the “striatal hyperdopaminergic” hypothesis. Of relevance, a recent animal study invoked elevated striatal dopamine as the driver of “hallucination-like percepts” in mice (Schmack et al., 2021). It will be important to confirm by physiology experiments if striatal dopamine signaling is enhanced in *Grin2a^+/-^* mutant mice, and whether any abnormal phenotypes (molecular or functional) can be rescued by D2 antipsychotic drugs. Also, given that Drd2, a bona fide drug target for SCZ, emerged as a highly changed gene from our snRNA-seq analysis of *Grin2a^+/-^* mice, we are hopeful that further study of transcriptomic changes in *Grin2a* mutants will uncover additional novel targets for SCZ treatment.

In summary, we show that loss of function of one copy of *Grin2a* in mice, analogous to rare *GRIN2A* mutations that cause SCZ in humans, affects EEG and behavior, and has impacts across widespread brain regions, diverse neuronal and non-neuronal cells, and multiple molecular pathways and processes in the juvenile and young adult mouse brain. The effects of *Grin2a* loss are not limited to glutamatergic synaptic signaling but involve other neurotransmitters and neuromodulators, including dopamine, serotonin, acetylcholine, and neuropeptides. Contextualizing these findings with human patient data highlights the *Grin2a* heterozygous mutant mouse as a new and compelling genetic model of SCZ and a promising resource for mechanistic enquiry and therapeutics discovery.

---

**Figure S1.**
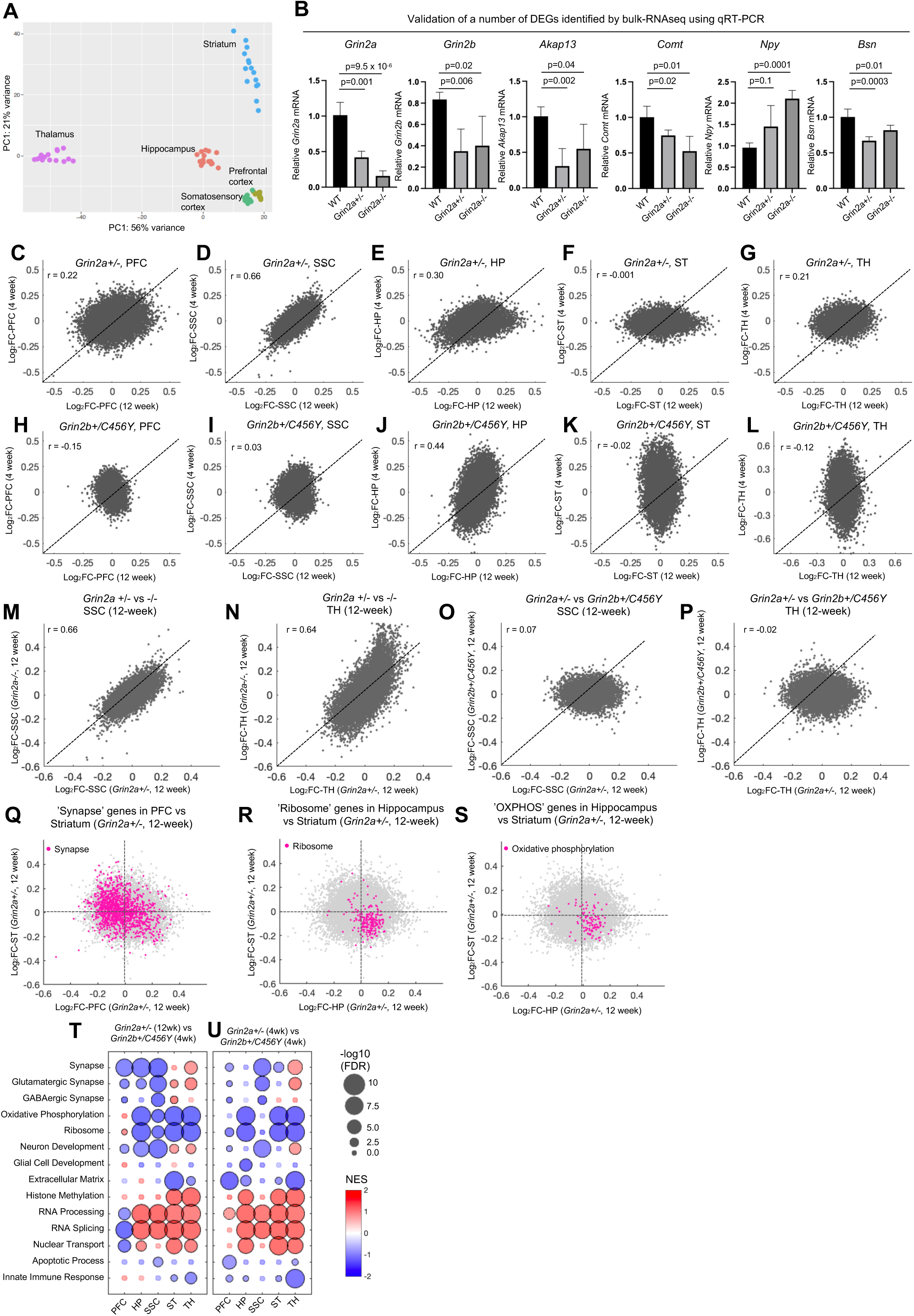
Bulk-RNA-seq analysis of five brain regions in *Grin2a* mutant mice, Related to Figure 1 and 2. (A) PCA plot showing the clustering of brain regions from 4-week *Grin2a* mutant animals and their wild-type littermates. Each circle represents one animal (n= 5 per genotype). PC, principal component. (B) Comparison of relative mRNA level measured by qRT-PCR (normalized to *Gapdh* mRNA) for several DEGs identified in the hippocampus of 12-week *Grin2a^+/-^* animals. Data are shown as mean +/- standard error (n=4-5 biological replicates). WT, wild type; p values are computed using two-tailed Student’s t test. (**C-L**) Gene expression correlation at 12 weeks versus 4 weeks in five tested brain regions in *Grin2a^+/-^* (**C-G**), and in *Grin2b^+/C456Y^* (**H-L**). Pearson’s r correlation values are indicated on the plots. (**M-P**) Gene expression correlation in the SSC (**M, O**) and thalamus (**N, P**) in *Grin2a^+/-^* versus *Grin2a^-/-^* (**M-N**), and in *Grin2a^+/-^* versus *Grin2b^+/C456Y^* (**O-P**) at 12 weeks. Pearson’s r correlation values are indicated on the plots. (**Q-S**) Transcriptomic changes in the PFC (**Q**) and hippocampus (**R-S**) versus the striatum in 12- week *Grin2a^+/-^* highlighting genes in the ‘Synapse’ (**Q**), ‘Ribosome’ (**R**) and ‘Oxidative phosphorylation’ (OXPHOS) (**S**) GO terms. (**T-U**) GSEA results for the studied brain regions using DE results from *Grin2a^+/-^* (12-week) versus *Grin2b^+/C456Y^* (4-week) comparison (**T**) and from *Grin2a^+/-^* (4-week) versus *Grin2b^+/C456Y^* (4-week) comparison (**U**), showing enrichment pattern of a selection of GO terms from MSigDB. Circles with black outlines indicate significance (FDR < 0.05).

**Figure S2.**
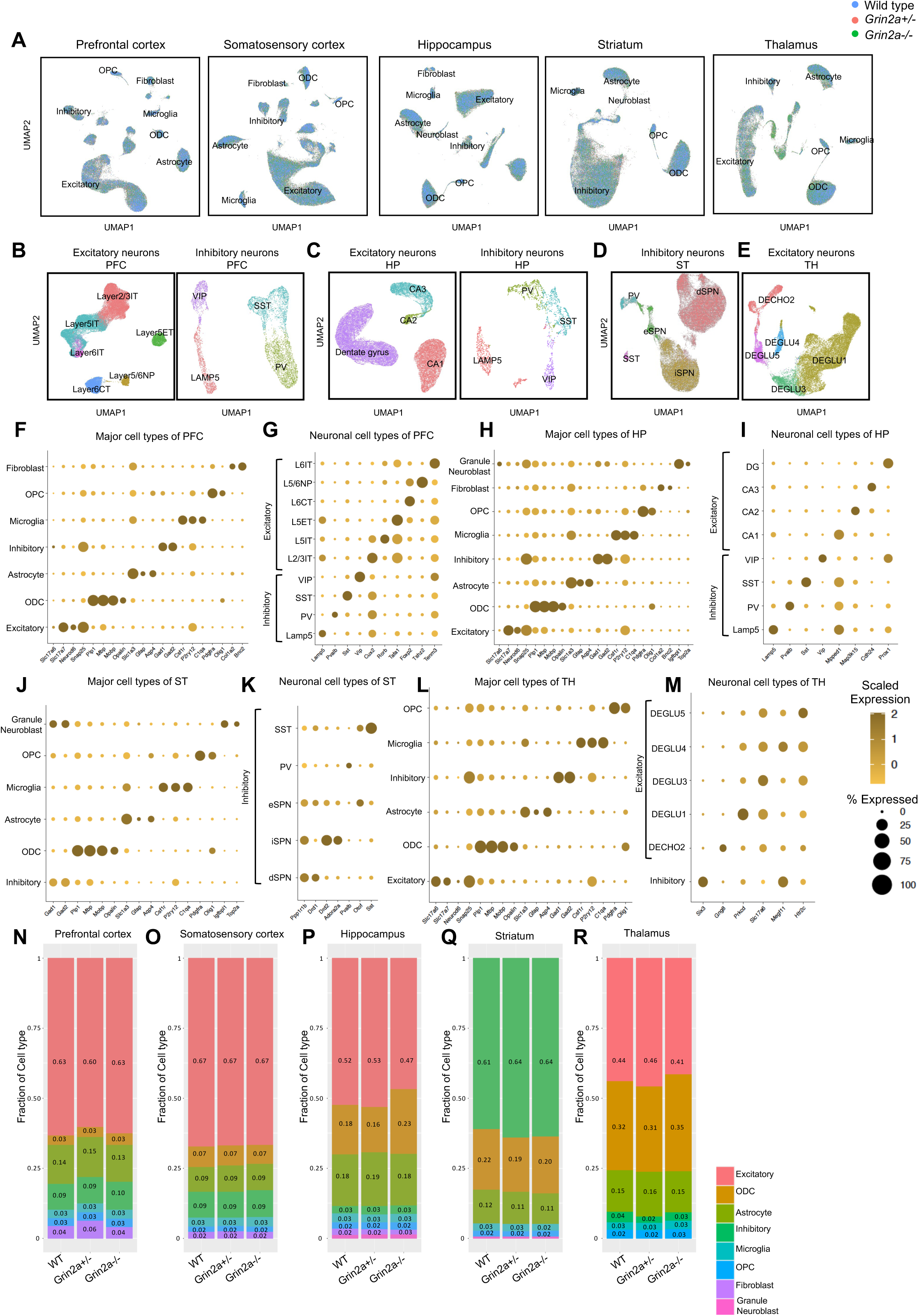
snRNA-seq analysis of five brain regions in *Grin2a* mutant mice, Related to Figure 3. (**A**) UMAP representation of major cell types identified in five brain regions in 12-week *Grin2a* animals. Nuclei are colored based on their genotype. (**B-E**) UMAP representation of excitatory and inhibitory neuron subtypes identified in the PFC (similar neuronal subtypes were identified in the SSC) (**B**), hippocampus (**C**), striatum (**D**) and thalamus (**E**) in 12-week *Grin2a* animals. Similar subtypes were identified in the PFC and hippocampus in 4-week *Grin2a* animals. (**F-M**) Dot plot showing the scaled expression (measured in transcripts per 10K over all nuclei) and percent of cells expressing marker genes of each cell type identified in the PFC (**F-G**), hippocampus (**H-I**), striatum (**J-K**) and thalamus (**L-M**) in 12-week *Grin2a* animals. Similar marker genes as in **F-I** were used to identify different cell types in 12-week SSC and 4-week PFC and in 4-week hippocampus datasets. (**N-R**) Fraction of each identified cell type in the PFC (**N**), SSC (**O**), hippocampus (**P**), striatum (**Q**) and thalamus (**R**) in 12-week *Grin2a* animals. Each column represents the average values for one genotype (n=4-6 animals per genotype). No significant difference was observed in the proportion of identified cell types between *Grin2a* mutant mice and their wild-type littermates (see Methods).

**Figure S3.**
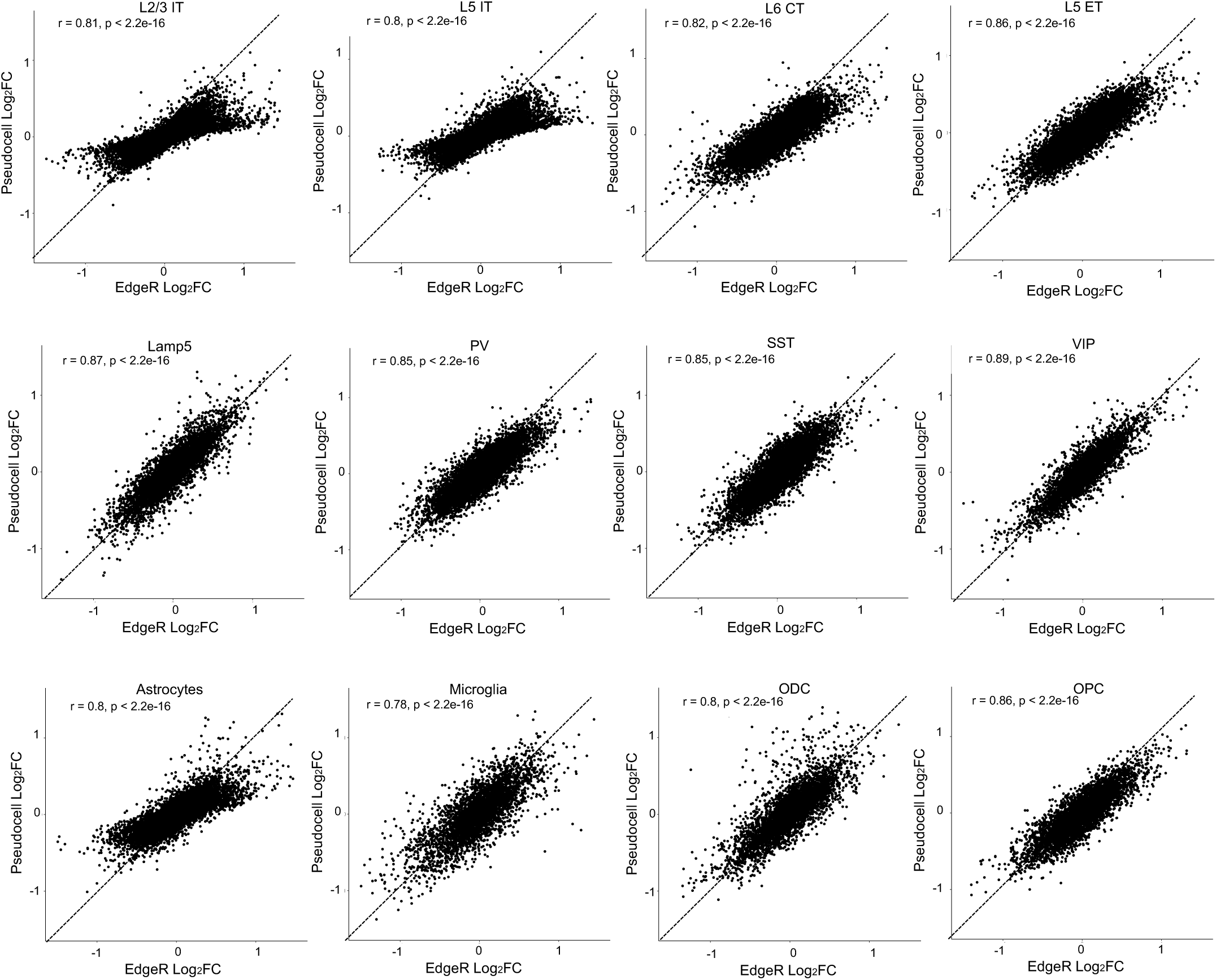
Gene expression changes are well correlated between pseudobulk and pseudocell analysis. Correlation between Log2FC values obtained from pseudobulk and pseudocell analysis across cell types in the PFC of 12-week *Grin2a^+/-^*. Pearson’s r and p values are indicated on the plot. Similar results were obtained for other brain regions, ages, and genotype.

**Figure S4.**
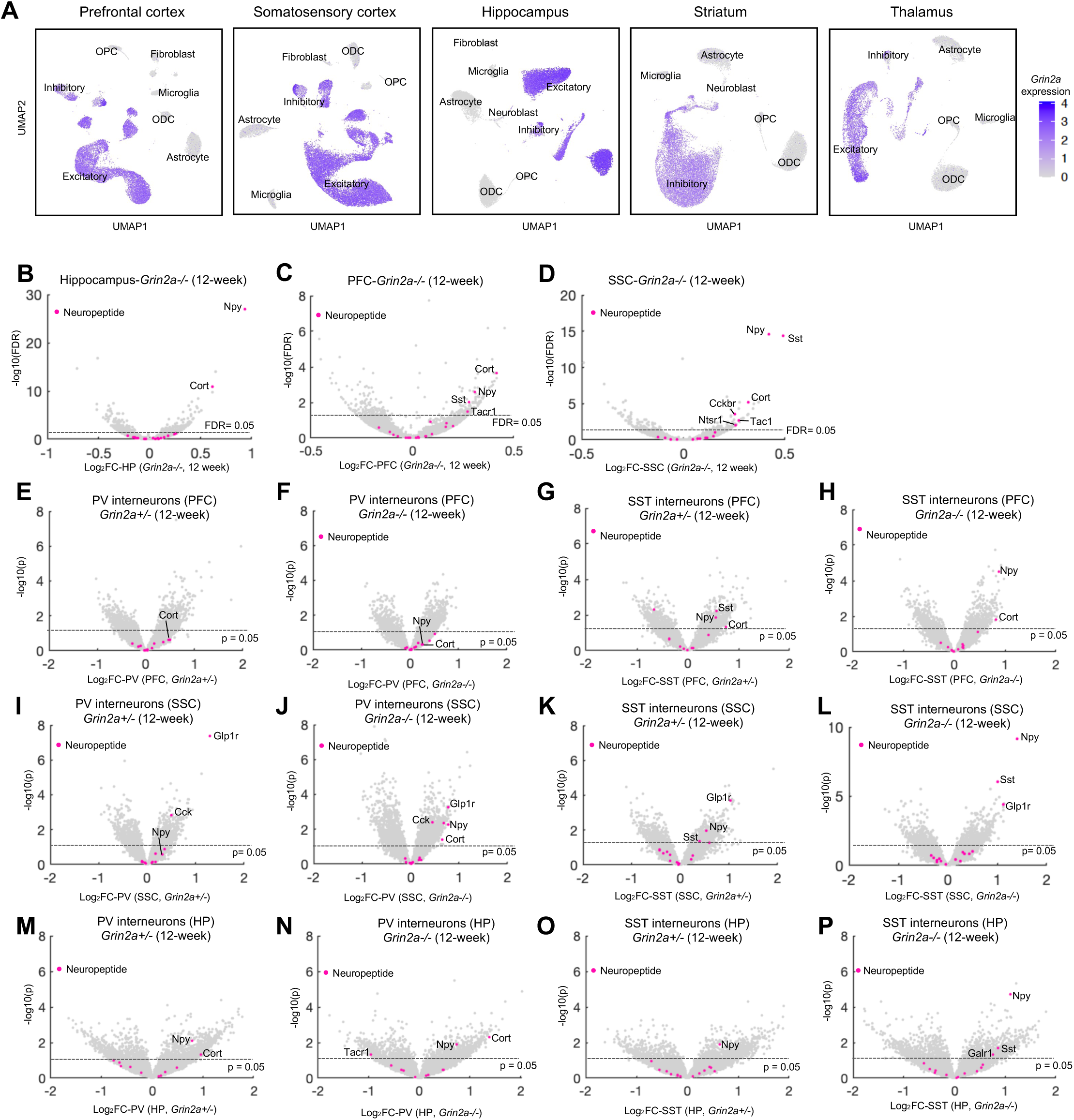
Changes in neuropeptide genes in inhibitory neurons of cortex and hippocampus in 12-week *Grin2a* mutants, Related to Figure 3. (**A**) UMAP representation of the major cell types identified in five brain regions in 12-week *Grin2a* animals. Nuclei are colored based on *Grin2a* expression. (**B-D**) Volcano plot of transcriptomic changes in bulk RNA-seq data from hippocampus (**B**), PFC (**C**), and SSC (**D**) in 12-week *Grin2a^-/-^* highlighting genes from the ‘neuropeptide’ gene-set (Table S6). (**E-P**) Volcano plot of transcriptomic changes in PV (**E-F**) and SST (**G-H**) interneuron of PFC, PV (**I-J**) and SST (**K-L**) interneuron of SSC, and PV (**M-N**) and SST (**O-P**) interneuron of hippocampus in 12-week *Grin2a^+/-^* and *Grin2a^-/-^* highlighting genes from the ‘neuropeptide’ gene-set (Table S6).

**Figure S5.**
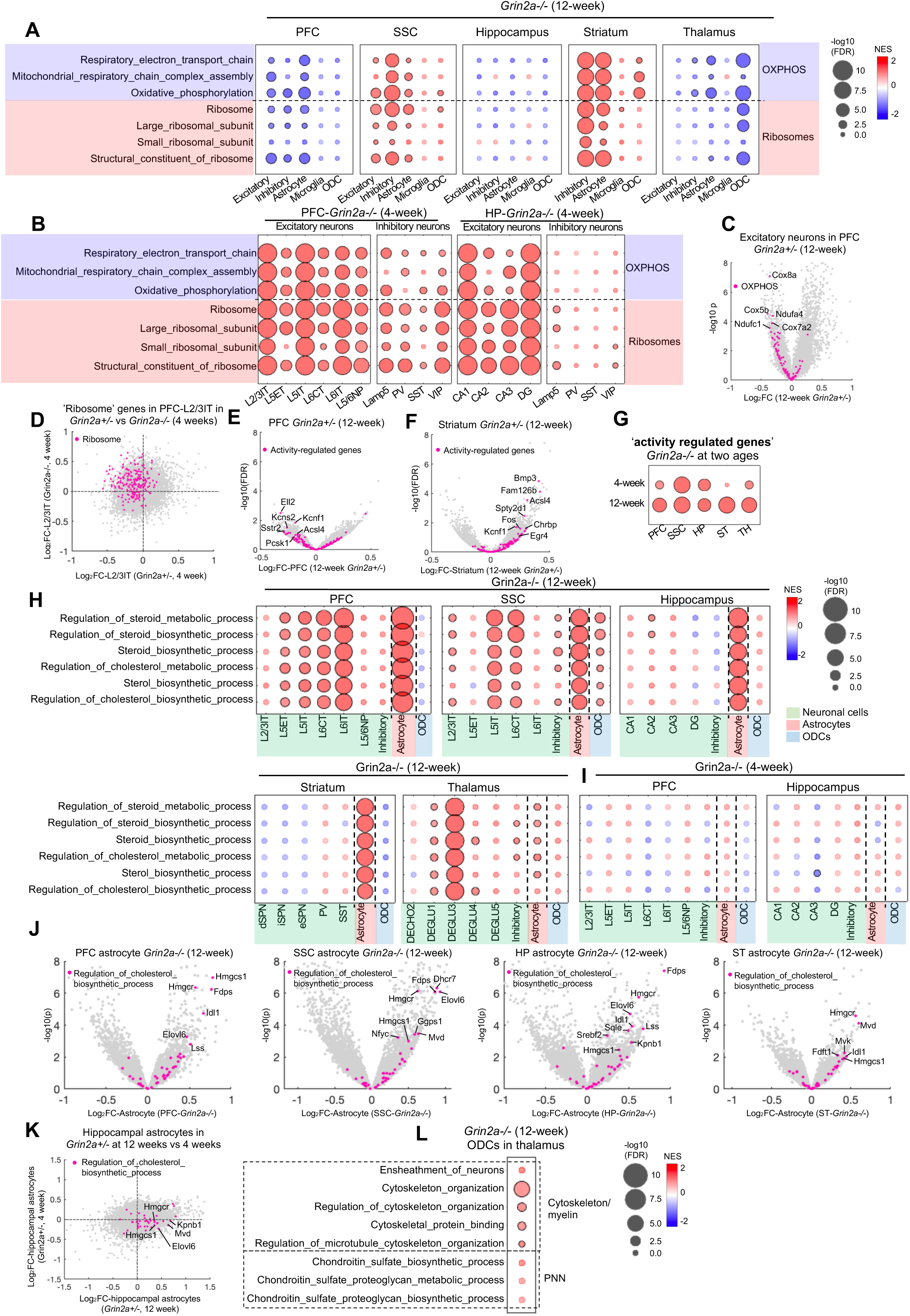
Changes of GO terms in neuronal and non-neuronal cells across brain regions in 4- and 12-week *Grin2a^-/-^* mutant mice, Related to Figure 4. (**A-B**) GSEA results from the 5 major cell types across brain regions in 12-week (**A**) and neuronal subtypes in 4-week (**B**) *Grin2a^-/-^* mutant mice. (C) Volcano plot of transcriptomic changes in excitatory neurons of PFC in 12-week *Grin2a^+/-^* highlighting genes from the ‘Oxidative phosphorylation’ (OXPHOS) GO term. (D) Transcriptomic changes in layer 2/3IT excitatory neurons of PFC in 4-week *Grin2a^+/-^* versus *Grin2a^-/-^* highlighting genes in the ‘Ribosome’ GO terms. (**E-F**) Volcano plot of transcriptomic changes in bulk RNA-seq data from the PFC (**E**) and striatum (F)) in 12-week *Grin2a^+/-^* highlighting genes from the ‘activity regulated’ gene-set (Table S6). (G) Enrichment of activity-regulated gene-set in the whole tissue transcriptome of the studied brain regions in *Grin2a^-/-^* at 4 and 12 weeks. (**H-I**) GSEA results using GO terms related to cholesterol biosynthesis in excitatory subtypes, inhibitory neurons, astrocytes and oligodendrocytes (ODCs) of the studied brain regions in 12- week (**H**) and 4-week (**I**) *Grin2a^-/-^* mutant mice. (J) Volcano plot of transcriptomic changes in astrocytes of PFC, SSC, hippocampus and striatum in 12-week *Grin2a^-/-^* highlighting genes from the ‘Regulation of cholesterol biosynthesis process’ GO term. (K) Transcriptomic changes in hippocampal astrocytes in *Grin2a^+/-^* at 12 weeks versus *Grin2a^+/-^* at 4 weeks highlighting genes from the ‘Regulation of cholesterol biosynthesis process’ GO term. (L) Changes in cytoskeleton/myelin and perineural net (PNN)-related GO terms in the oligodendrocytes of thalamus in 12-week *Grin2a^-/-^* mutant mice. In all the bubble plots circles with black outlines indicate significance (FDR < 0.05).

**Figure S6.**
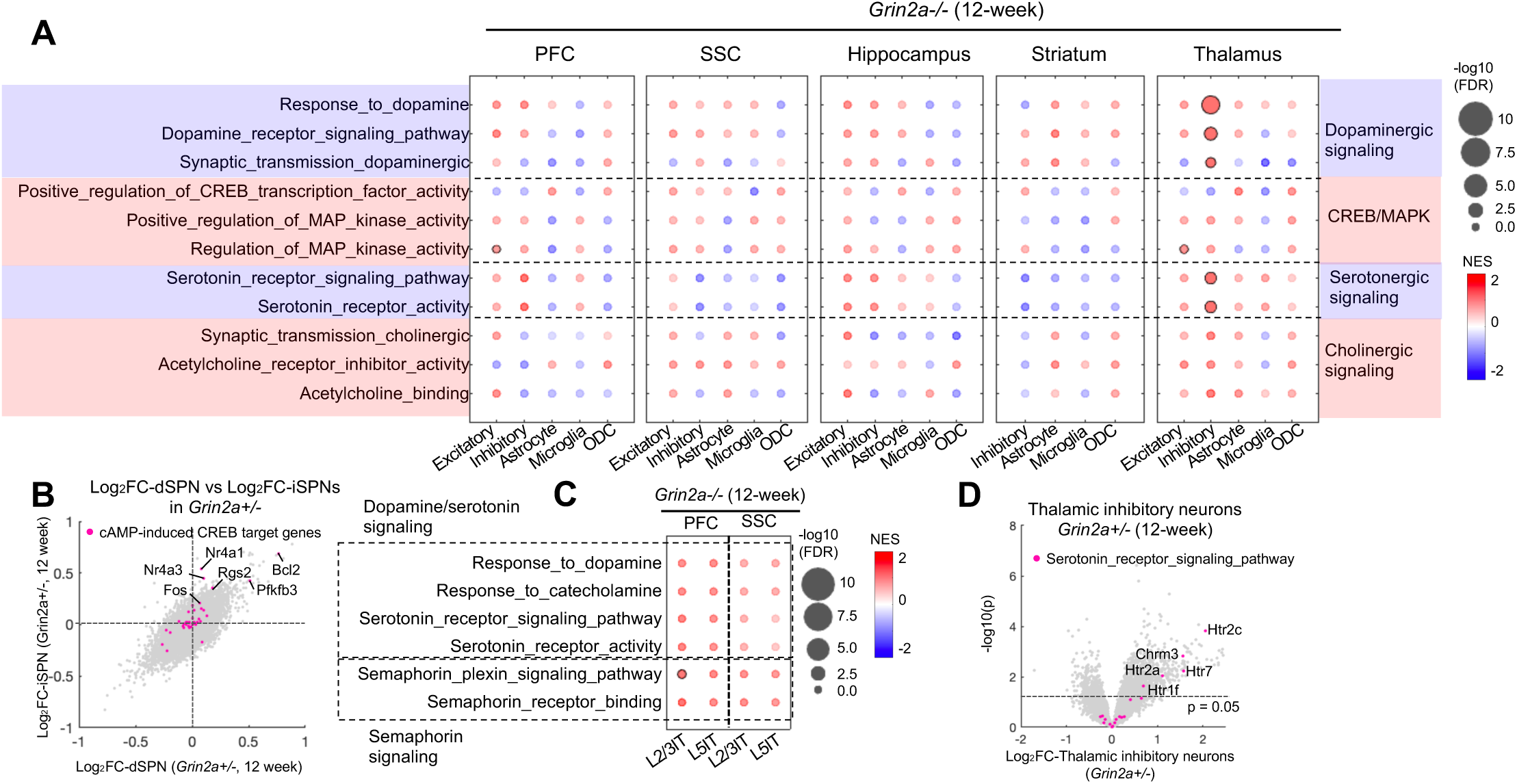
Changes in dopamine signaling in the striatum of *Grin2a* mutant mice, Related to Figure 5. (A) GSEA results from the 5 major cell types across brain regions in 12-week *Grin2a^-/-^* mutant mice. (B) Transcriptomic changes in dSPNs versus iSPNs in the striatum of 12-week *Grin2a^+/-^* highlighting the cAMP-induced CREB target genes (Table S6). (C) Changes in serotonin, dopamine and semaphorin signaling GO terms in layer 2/3IT and layer 5IT in PFC and SSC at 12 weeks in *Grin2a^-/-^*. (D) Volcano plot of transcriptomic changes in inhibitory neurons of thalamus in 12-week *Grin2a^+/-^* highlighting genes from the ‘Serotonin receptor signaling pathway’ GO term.

**Figure S7.**
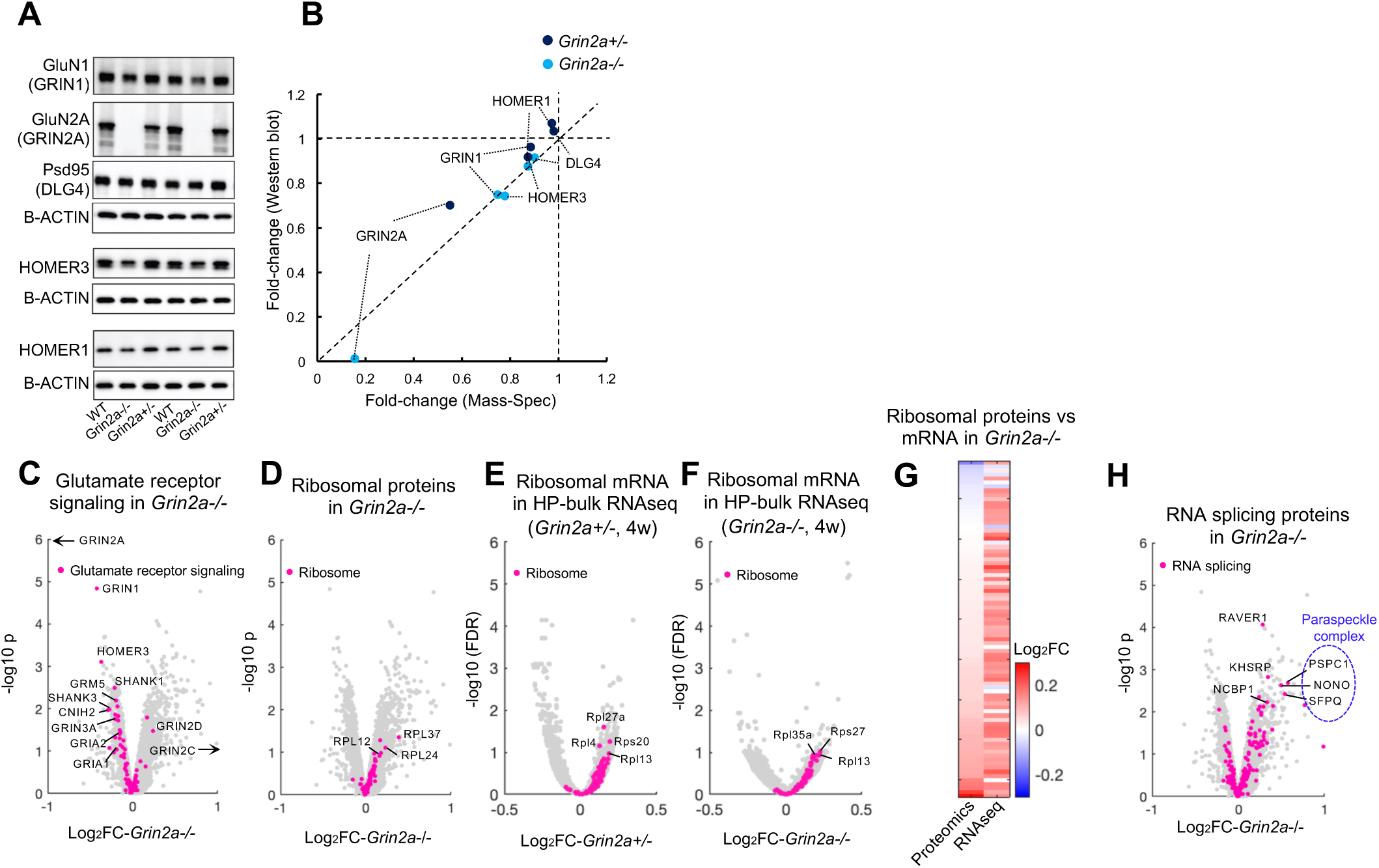
Synapse proteome changes in *Grin2a* mutant mice, Related to Figure 6. (A) Western blots probing for GluN1, GluN2A, Psd95, HOMER3 and HOMER1 in synaptic fractions from an independent cohort of *Grin2a* mutant mice. (B) Quantification of (**A**) against fold-changes obtained from proteomics data. (C) Volcano plot of proteomic changes in *Grin2a^-/-^* synapses highlighting proteins in the ‘glutamate receptor signaling pathway’ gene-set. (D) Volcano plot of proteomic changes in *Grin2a^-/-^* synapses highlighting proteins in the ‘cytosolic ribosome’ gene-set. (**E-F**) Volcano plot of the whole tissue transcriptomic changes in the hippocampus of *Grin2a^+/-^* (**E**) and *Grin2a^-/-^* (**F**) at 4 weeks highlighting proteins in the ‘cytosolic ribosome’ gene-set. (G) Heatmap depicting all detected proteins in the ‘cytosolic ribosome’ gene-set in the hippocampal synaptic proteome and their mRNA changes in hippocampal bulk RNA-seq data in 4-week *Grin2a^-/-^*. (H) Volcano plot of proteomic changes in *Grin2a^-/-^* synapses highlighting proteins in the ‘RNA splicing’ gene-set. In **C** arrows indicate the proteins that are out of the range of the plots.

**Figure S8.**
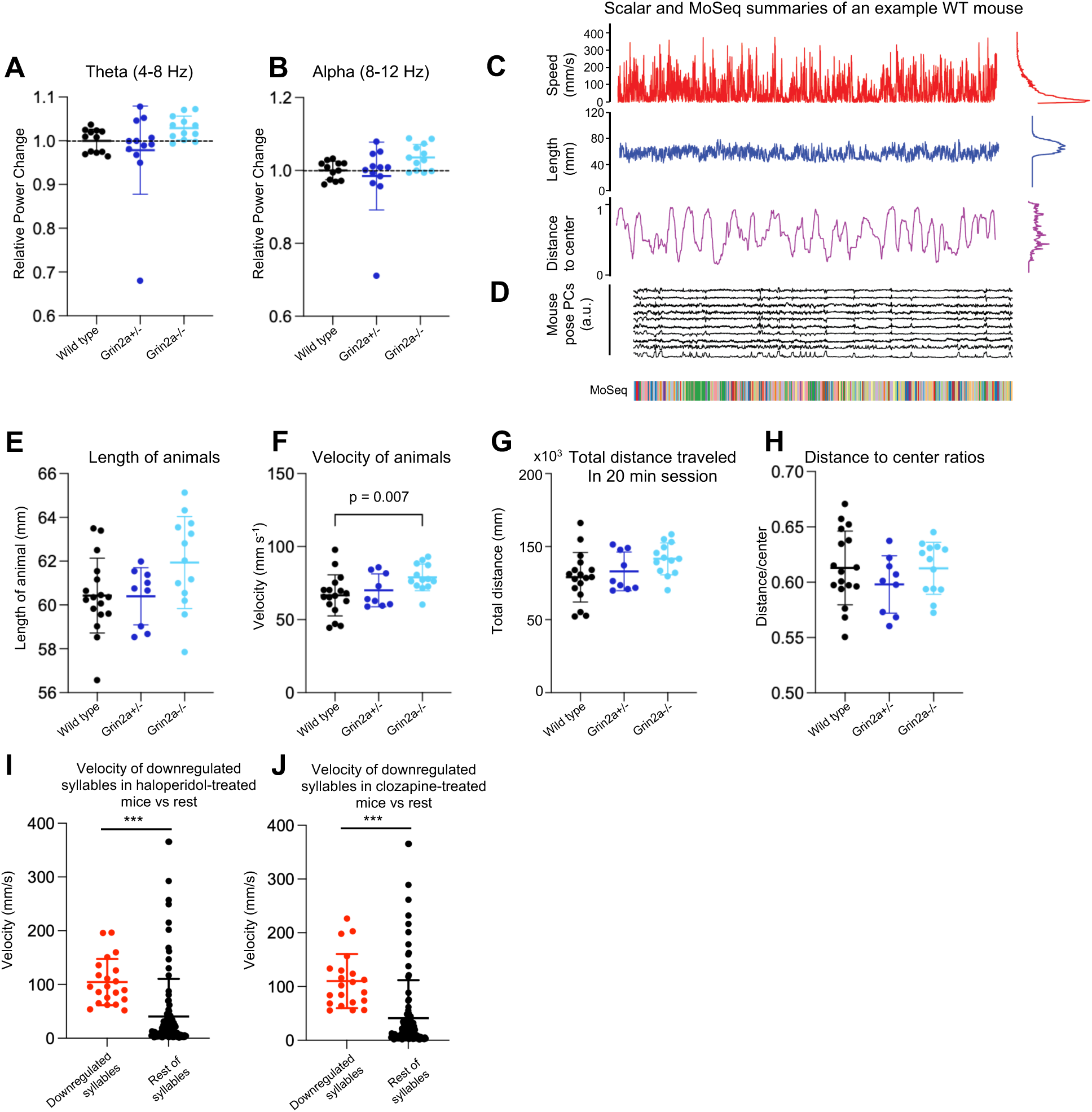
EEG and MoSeq characterization of *Grin2a* mutant mice, Related to Figure 7. (**A-B**) Power spectral density (PSD) changes in GRIN2A mutant mice relative to wild-type animals at ∼3 months of age, measured during NREM sleep during the light cycle for each of the frequency bands: (**A**) theta, 4-8 Hz; (**B**) alpha, 8-12 Hz gamma. Error bars show mean +/- standard error; n=12 mice/group. (C) Temporal distribution of speed (mm/s), mouse length (mm), and distance-to-center ratio (with 0 representing the center and 1 representing the absolute margin of arena) for a representative wild-type mouse. (D) The top 10 principal component (PC) traces of the processed 3D depth images for an example wild-type mouse. The MoSeq Autoregressive Hidden Markov Model (AR-HMM) (see Methods) takes in the top 10 PC values as inputs and segments the depth videos into modulated behavioral motifs, referred to as syllables. For each mouse, a MoSeq-based behavioral summary was generated using 20 min of data (bottom). (**E-H**) Length (**E**), velocity (**F**), total distance traveled within 20 min session (**G**), and distance to center ratio (**H**) are plotted per genotype. Error bars show mean +/- standard error; n=17 wild type (WT), n=9 *Grin2a^+/-^*, n=13 *Grin2a^-/-^*. p values are computed using Welch’s t test. (**I-J**) Mean velocity for downregulated syllables in haloperidol- (**I**) and clozapine- (**J**) treated mice (red box in Fig. 7I) versus rest of identified syllables. Error bars show mean +/- standard error. Asterisks (*) indicate statistical significance assessed using non-parametric permutation tests (see Methods); (***) indicates p < 0.001.

## LIST OF SUPPLEMENTAL TABLES

**Table S1.** DEGs in bulk RNA-seq for all brain regions and ages in *Grin2a^+/-^* and *Grin2a^-/-^*, Related to Figure 1.

**Table S2.** DEGs in bulk RNA-seq for all brain regions and ages in *Grin2b^+/C456Y^*, Related to Figure 1.

**Table S3.** Number of reads per sample for bulk RNA-seq analysis including all brain regions, ages, and genotypes, Related to Figure 1.

**Table S4.** GSEA results for all brain regions and ages in *Grin2a^+/-^* and *Grin2a^-/-^*, Related to Figure 2.

**Table S5.** GSEA results for all brain regions and ages in *Grin2b^+/C456Y^*, Related to Figure 2.

**Table S6.** List of curated gene sets.

**Table S7.** List of Marker genes used for cell type annotations and alignment statistics as well as QC metrics per snRNA-seq sample, Related to Figure 3.

**Table S8.** DEGs identified by pseudobulk approach in all cell types of 4 week-PFC.

**Table S9.** DEGs identified by pseudobulk approach in all cell types of 12 week-PFC.

**Table S10.** DEGs identified by pseudobulk approach in all cell types of 4 week-Hippocampus.

**Table S11.** DEGs identified by pseudobulk approach in all cell types of 12 week-Hippocampus.

**Table S12.** DEGs identified by pseudobulk approach in all cell types of 12 week-SSC.

**Table S13.** DEGs identified by pseudobulk approach in all cell types of 12 week-Striatum.

**Table S14.** DEGs identified by pseudobulk approach in all cell types of 12 week-Thalamus.

**Table S15.** GSEA results for the pseudobulk approach in all cell types of 4 week-PFC.

**Table S16.** GSEA results for the pseudobulk approach in all cell types of 12 week-PFC.

**Table S17.** GSEA results for the pseudobulk approach in all cell types of 4 week-Hippocampus.

**Table S18.** GSEA results for the pseudobulk approach in all cell types of 12 week-Hippocampus.

**Table S19.** GSEA results for the pseudobulk approach in all cell types of 12 week-SSC.

**Table S20.** GSEA results for the pseudobulk approach in all cell types of 12 week-Striatum.

**Table S21.** GSEA results for the pseudobulk approach in all cell types of 12 week-Thalamus.

**Table S22.** List of number of nuclei per cell type per sample and DEGs identified by pseudocell approach in all cell types of 4 week-PFC.

**Table S23.** List of number of nuclei per cell type per sample and DEGs identified by pseudocell approach in all cell types of 12 week-PFC.

**Table S24.** List of number of nuclei per cell type per sample and DEGs identified by pseudocell approach in all cell types of 4 week-Hippocampus.

**Table S25.** List of number of nuclei per cell type per sample and DEGs identified by pseudocell approach in all cell types of 12 week-Hippocampus.

**Table S26.** List of number of nuclei per cell type per sample and DEGs identified by pseudocell approach in all cell types of 12 week-SSC.

**Table S27.** List of number of nuclei per cell type per sample and DEGs identified by pseudocell approach in all cell types of 12 week-Striatum.

**Table S28.** List of number of nuclei per cell type per sample and DEGs identified by pseudocell approach in all cell types of 12 week-Thalamus.

**Table S29.** GSEA results for the pseudocell approach in all cell types of 4 week-PFC.

**Table S30.** GSEA results for the pseudocell approach in all cell types of 12 week-PFC.

**Table S31.** GSEA results for the pseudocell approach in all cell types of 4 week-Hippocampus.

**Table S32.** GSEA results for the pseudocell approach in all cell types of 12 week-Hippocampus.

**Table S33.** GSEA results for the pseudocell approach in all cell types of 12 week-SSC.

**Table S34.** GSEA results for the pseudocell approach in all cell types of 12 week-Striatum.

**Table S35.** GSEA results for the pseudocell approach in all cell types of 12 week-Thalamus.

**Table S36.** DEPs identified in synaptic proteome of 4 week-Hippocampus, Related to Figure 6.

**Table S37.** GSEA results for synaptic proteome of 4 week-Hippocampus, Related to Figure 6.

## METHODS

### *Grin2a* and *Grin2b* mutant mice

*Grin2a*^-/-^ mice, originally generated as described (Kadotani et al., 1996), were obtained from RIKEN BioResource Research Center (RBRC02256) and were crossed against wild-type C57/BL6J mice (Jackson Laboratory, #000664) to generate *Grin2a*^+/-^ mice. The *Grin2a^+/-^* mice were then crossed with each other to produce *Grin2a^+/-^*, *Grin2a*^-/-^ and wild-type littermates used in all the experiments. *Grin2a* mutant mice were housed at AAALAC-approved facilities on a 12- hour light/dark cycle, with food and water available *ad libitum*. All procedures involving *Grin2a* mutant mice were approved by the Broad Institute IACUC (Institutional Animal Care and Use Committee) and conducted in accordance with the NIH Guide for the Care and Use of Laboratory Animals.

Frozen whole brain tissues of *Grin2b*^+/C456^ and their wild-type littermates, all under the genetic background of C57BL/6J, were kindly provided by Dr. Eunjoon Kim (Department of Biological Sciences, Korea Advanced Institute of Science and Technology, South Korea).

### Brain perfusion and dissection

Brain tissues for bulk and snRNA-seq were prepared as described at protocols.io (https://www.protocols.io/view/fresh-frozen-mouse-brain-preparation-for-single-nu-bcbrism6). To minimize one source of variability, only male mice were used in this study. Briefly, at 2, 4, 12 and 20 weeks of age, male *Grin2a*^+/-^, *Grin2a*^-/-^ and wild-type mice were anesthetized by administration of isoflurane in a gas chamber. While anesthesia was prolonged via a nose cone through which 3% isoflurane flowed, transcardial perfusions were performed with ice-cold Hank’s Balanced Salt Solution (HBSS) to remove blood from the brain. The brains were immediately frozen for three minutes in liquid nitrogen vapor and stored at –80 °C until dissection.

Brain dissection was performed in the cryostat (Leica CM3050S), and all the tools required for dissection were precooled to –20 °C. The cerebellum was first removed with a cut in the coronal plane using a razor blade precooled to –20 °C. The brains were then cut at the midsagittal plane to use the right and left hemispheres of each brain for bulk and snRNA-seq experiments, respectively. To dissect the prefrontal cortex, each brain hemisphere was securely mounted from lateral surface onto a cryostat chuck with O.C.T. (optimal cutting temperature) freezing medium (Tissue-Tek) such that the midsagittal surface was left exposed and thermally unperturbed. The medial prefrontal cortex was then dissected by hand in the cryostat using an ophthalmic microscalpel (Feather safety Razor no. P-715) precooled to –20 °C. To dissect the other brain regions, the remaining brain hemisphere was removed from the chuck and mounted from the olfactory bulb onto a new cryostat chuck with O.C.T. freezing medium. Coronal sections were cut by advancing the cryostat 30 μm at a time in trimming mode until the dorsal hippocampus was visible. This stepwise approach reduced disruption of the brain tissue surface that could occur with larger steps. The dorsal hippocampus was then dissected by hand using the ophthalmic microscalpel, and the thalamus was collected using a precooled biopsy punch (Thermo Fisher Scientific) with 0.15 cm diameter. 30 μm-coronal sections were further cut by advancing the cryostat until the lateral ventricles became triangle-shaped. The somatosensory cortex was then dissected by hand using the ophthalmic microscalpel and the dorsal striatum was collected using a precooled biopsy punch with 0.15 cm diameter. All the brain regions were identified according to the Allen Brain Atlas. Each excised tissue was placed into a precooled 1.5-ml PCR tube using precooled forceps and stored at –80°C.

### RNA isolation and library preparation for bulk RNA-seq analysis

RNA was prepared from microdissected tissues using RNeasy Mini Kit (Qiagen) following the manufacturer’s instructions. Briefly, tissue samples were lysed and homogenized by hand, placed into columns, and bound to the RNeasy silica membrane. Next, contaminants were washed away, columns were treated with DNase (Qiagen) to digest residual DNA, and concentrated RNA was eluted in water. RNA concentration was measured using a NanoDrop Spectrophotometer and RNA integrity (RIN) was measured with RNA pico chips (Agilent) using a 2100 Bioanalyzer Instrument (Agilent). Purified RNA was then stored at -80°C until bulk sequencing library preparation.

Bulk sequencing libraries were prepared using a TruSeq Stranded mRNA Kit (Illumina) following the manufacturer’s instructions. 200 ng of isolated total RNA from each sample was used and the concentration of resulting cDNA library was measured with High Sensitivity DNA chips (Agilent) using a 2100 Bioanalyzer Instrument (Agilent). A 10nM normalized library was pooled, and sequencing was performed on a NovaSeq S2 (Illumina) with 50 bases each for reads 1 and 2 and 8 bases each for index reads 1 and 2.

### qRT-PCR

RNA was extracted with the RNeasy Mini Kit (Qiagen) as stated above. qRT-PCR was performed using the TaqMan RNA-to-Ct 1-Step Kit and TaqMan gene expression assays (Thermo Fisher Scientific). PCR amplifications were performed in triplicate or quadruplicates (n = 4-5 animals per genotype) and gene expression was determined by the comparative cycle threshold (ΔΔCt) method using *Actb* or *Gapdh* as an internal housekeeping gene control.

### Nuclei extraction and library preparation for snRNA-seq analysis

Brain tissue dissection and nuclei extraction were performed on the same day to avoid freeze-thaw cycles which would result in poor quality nuclei. A gentle, detergent-based dissociation was used to extract the nuclei, according to a previously published protocol (Biancalani et al., 2021), also available at protocols.io (https://www.protocols.io/view/frozen-tissue-nuclei-extraction-bbseinbe). Extracted nuclei were then loaded into the 10x Chromium V3.1 system (10x Genomics) and library preparation was performed according to the manufacturer’s protocol. A 10nM normalized library was pooled, and sequencing was performed on a NovaSeq S2 (Illumina) with 28 and 75 bases for reads 1 and 2 and 10 bases each for index reads 1 and 2.

### Isolation of synaptic fractions and preparation for Mass Spectrometry

P28-32 *Grin2a* mutant mice were sacrificed using CO2 anesthesia, after which hippocampi were dissected rapidly, flash-frozen on dry ice, and stored at -80°C. Synapse fractions were purified as described previously (Dejanovic et al., 2018; Dejanovic et al., 2022). Briefly, hippocampi were thawed and dounce-homogenized (10 strokes) in ice-cold homogenization buffer (5 mM HEPES pH 7.4, 1 mM MgCl2, 0.5 mM CaCl2, supplemented with phosphatase (Sigma 4906837001) and protease inhibitors (Sigma 4693159001)). The homogenate was centrifuged for 10 minutes at 1,400 g (4°C) and the resulting supernatant was re-centrifuged at 13,800 g for 10 minutes (4°C). The resulting pellet was resuspended in 0.32 M Sucrose, 6 mM Tris-HCl (pH 7.5) and layered gently on a 0.85 M, 1 M, 1.2 M discontinuous sucrose gradient (all layers in 6mM Tris-HCl pH 7.5) and ultracentrifuged at 82,500 g for 2 hours (4°C). The synaptosome fraction which sediments at the 1M and 1.2M sucrose interface, was collected, an equal volume of ice-cold 1% Triton X-100 (in 6mM Tris-HCl pH 7.5) was added, mixed thoroughly, and incubated on ice for 15 minutes. The mixture was ultracentrifuged at 32,800 g for 20 minutes (4°C), and the pellet (the PSD fraction) was collected by resuspension in 1% SDS. A small aliquot was taken to measure the protein concentration, and the remaining protein was stored at -80°C until being processed for mass spectrometry or Western blotting / SDS-PAGE.

Sixteen synaptic fractions extracted from the hippocampus of wild type, *Grin2a*^+/-^ and *Grin2a*^-/-^ mouse brains were used for the proteomics study using tandem mass tag (TMT) isobaric labeling strategy for quantitation (n=5 wild type; n=6 Grin2a^+/-^; n=5 Grin2a^-/-^). All samples contained 35 ug protein pre-digestion and were processed using the following workflow.

Samples were prepared in a one-percent SDS buffer to ensure proper dissolution of membrane proteins and digested using single-pot, solid-phase-enhanced sample-preparation (SP3) technology. Reduction of proteins using 5 mM dithiothreitol and alkylation of them using 10 mM iodoacetamide was performed at 60°C and room temperature respectively. All samples were diluted to a total volume of 140 uL with a working solution consisting of: 50 mM HEPES, 5mM EDTA, 50 mM NaCl, 2 ug/mL aprotinin, 10 ug/mL leupeptin, 1 mM PMSF, 10 mM NaF, 1:100 PIC2 and PIC3 and 1% SDS. The proteins were then bound to SP3 paramagnetic beads in a 1:10 protein to bead ratio using 100% ethanol to induce binding. Samples were then incubated at 24°C for 5 minutes mixing at 1000 rpm and then placed on a magnetic rack to pull down the beads. Beads were kept immobilized and washed three times with 1 mL of 80% ethanol, then resuspended in 100 mM ammonium bicarbonate to maintain a 1 ug/uL bead concentration. Each sample was sonicated in a water bath for 1 minute and gently vortexed to homogenize. Then sequential digestion steps were performed using 1:25 enzyme to substrate ratio of Lys-C for 2 hours and Trypsin overnight at 37 °C. Following digestion, samples were centrifuged at 12,000 rpm for 5 minutes then placed on a magnetic rack where the supernatant was collected and acidified to a 1% final concentration of formic acid.

Post-digest peptide measurements were performed by Nanodrop Spectrophotometer (Thermo Fisher Scientific) and 30 ug aliquots per sample were made for labeling. A TMT16 plex was constructed by randomly assigning the samples from each group to channels within the plex. Samples were reconstituted in 50 mM HEPES buffer for labeling and 20 uL of 25 ug/uL TMT16 reagent was added for the labeling reaction. After confirming successful labeling (>95% label incorporation), the reactions were quenched with 5% hydroxylamine and combined. The mixed sample was then desalted on a 50 mg tC18 SepPak cartridge and fractionated on a 2.1 mm x 250mm Zorbax 300 extend-c18 reverse-phased column. One-minute fractions were collected during the entire elution and then concatenated into 12 fractions.

One microgram of each proteome fraction was analyzed on a QE mass spectrometer (Thermo Fisher Scientific) coupled to an easy-nLC 1200 LC system (Thermo Fisher Scientific). Samples were separated using 0.1% Formic acid/3% Acetonitrile as buffer A and 0.1% Formic acid /90% Acetonitrile as buffer B on a 27cm 75um ID picofrit column packed in-house with Reprosil C18- AQ 1.9um beads (Dr Maisch GmbH) with a 90min gradient consisting of 6-20% B in 62min, 20- 30% B for 22min, 30-60% B in 9min, 60-90% B for 1min followed by a hold at 90% B for 5min. The MS method consisted of a full MS scan at 70,000 resolution and an AGC target of 3e6 from 300-1800 m/z followed by MS2 scans collected at 35,000 resolution with an AGC target of 5e4 with a maximum injection time of 120ms and a dynamic exclusion of 15 seconds. The isolation window used for MS2 acquisition was 0.7 m/z and 12 most abundant precursor ions were fragmented with a normalized collision energy (NCE) of 27 optimized for TMT16 data collection.

### Western blotting / SDS-PAGE

The protein concentrations of synaptic fractions or total brain lysate were determined using Bicinchoninic acid assay (BCA; Pierce 23227). To equalize protein concentrations, samples were diluted with 6X SDS-Sample buffer (Boston Bioproducts BP-111R; to a final concentration of 1X) and water. The diluted samples were then heated at 95°C for 5-7 minutes after which they were stored at -20°C until SDS-PAGE. Before running on a gel, samples were thawed, heated at 95°C for 1 minute and centrifuged briefly. Equal amounts of protein in an equal volume were loaded for each sample on 4-20% Tris-glycine polyacrylamide gels; the gels were run using Tris-glycine SDS running buffer at constant voltage. Proteins were transferred to 0.2 μm Nitrocellulose membranes using semi-dry transfer (Bio-Rad Transblot Turbo; 25V 30 minutes). Membranes were blocked using 5% milk in Tris-buffered saline supplemented with 0.1% Tween-20 (TBST) for 1.5 hours at room-temperature (RT). Membranes were then probed overnight (4 °C) with primary antibodies (1:1000 for all antibodies) in 5% milk TBST with gentle rotation. After four vigorous 5 -minute washes in TBST, membranes were incubated with either 1:5,000-10,000 HRP- or Biotin-conjugated (Jackson Immunoresearch) anti-IgG antibody raised against the appropriate species in 5% TBST for 90 minutes (RT). The latter membranes were washed four times (5 minutes each, RT) vigorously in TBST and labeled further with 1:10,000 Streptavidin Poly-HRP (Pierce 21140; in 5% milk TBST at RT, 1 hour). All membranes were then washed four times in TBST and imaged on the FluorChem-E (Protein Simple) or the ChemiDoc MP (Bio-Rad) platforms. Enhanced chemiluminescence and exposures were optimized to obtain signals in the linear range. Membranes were then stripped, re-blocked with 5% milk TBST, re-probed with a different primary antibody and imaged again as described.

1 μg protein of synaptic fraction samples were loaded in gels probing for HOMER1 and HOMER3, while 3.5 μg protein was loaded in those probing for GluN2A, GluN1, and Psd95. For COMT 20 μg protein total prefrontal or striatal lysates were loaded in gels. HOMER1, HOMER3, Psd95 and COMT were probed using HRP-conjugated anti-IgG antibodies, while GluN2A and GluN1 were probed using the Biotin-Streptavidin-poly HRP approach. HOMER1, HOMER3, and GluN1 were probed on fresh membranes while Psd95, GluN2A were probed on a membrane previously probed for GluN1. β-ACTIN was probed using an HRP-conjugated primary antibody subsequent to imaging of other proteins.

The details of the primary antibodies used, and species they were raised in, are as follows:

| Antibody | Source | Identifier | Species |
| --- | --- | --- | --- |
| COMT | BD Biosciences | Cat# 611970 | Mouse |
| GluN1 | Cell signaling | Cat# 5704 | Rabbit |
| GluN2A | Cell signaling | Cat# 4205 | Rabbit |
| HOMER1 | Synaptic systems | Cat# 160 003 | Rabbit |
| HOMER3 | Synaptic systems | Cat# 160 303 | Rabbit |
| Psd95 | BioLegend | Cat# 810401 | Mouse |
| β-ACTIN | Sigma Aldrich | Cat# A3854 | Mouse; HRP-conjugated |

### EEG implantation surgery and recording

6- to 9-week-old mice (n=12 mice/group) were deeply anesthetized with isoflurane. A prefabricated EEG/EMG headmount (#8201-SS, Pinnacle Technology, Lawrence, KS) was secured to the skull with three 0.10’’ intracranial electrode screws (#8403, Pinnacle Technology) at the following stereotactic coordinates: parietal recording electrode (-2 AP, 1.5 ML to Bregma), ground and reference electrodes (bilaterally -1 AP, 2 ML to Lambda). The electromyogram (EMG) electrodes were placed bilaterally in the nuchal muscles. Electrodes were soldered to the EEG/EMG head-mount and dental acrylic was used to secure the connections. Animals were given at least one week of post-operative recovery before EEG recording. Following recovery from EEG implantation, mice were tethered to the Pinnacle recording system, with at least 24 hours of habituation before recording. For sleep/wake recordings, EEG/EMG signals were recorded for 24 hours from the onset of the dark phase, ZT12 (6 pm EDT or 7 pm EST). Animals remained tethered to the Pinnacle system throughout the testing period with *ad libitum* access to food and water. All signals were digitized at a sampling rate of 1000 Hz, filtered (1–100 Hz bandpass for EEG; 10–1 kHz bandpass for EMG), and acquired using the Sirenia Acquisition program (Pinnacle Technology). EEG recordings took place at a timepoint roughly corresponding to 3-months of age (age range: 8-10 weeks).

### Motion Sequencing (MoSeq)

MoSeq open field assay (OFA) behavioral analysis was done on mouse 3D pose data recorded in the MoSeq recording apparatus. The apparatus included a black matte plastic bucket as the arena where the mouse ran and a rigid cage that enclosed the arena. A Kinect for Windows v.2 (Microsoft) depth sensing camera was stably mounted on the cage to record the mouse’s behavior in 3D from a top-down view. Animals were habituated to the behavioral room in their home cage for 10 minutes before recording, and all MoSeq sessions were conducted during the mouse’s dark cycle in a red light-lit room. Each session was recorded for 20 minutes at a sampling rate of 30 frames per second. Before and between each recording session, the arena was wiped down with water, then 70% ethanol, then water, and then wiped dry. Experimental metadata such as Session Name and Subject names were saved along with the depth videos. 16 wild-type, 9 *Grin2a*^+/-^ and 13 *Grin2a*^-/-^ mice were used for MoSeq OFA recordings.

## COMPUTATIONAL METHODS

### Bulk RNA-seq data analysis

RSEM v1.3.0 (Li and Dewey, 2011) was used to estimate gene and isoform level expression values for our bulk RNA-seq samples. The M25 (GRCm38.p6) GENCODE reference was generated using RSEM-prepare-reference with default parameters. Expression values were calculated using RSEM-calculate-expression with the following flags: --bowtie2, --paired-end, -- estimate-rspd, --append-names, and --sort-bam-by-coordinate. The DESeq2 v1.20.0 (Love et al., 2014) package was used to run DE analysis of each of the brain regions and ages. All replicates for a given age and brain region were read into a single DESeq object to ensure that normalization was consistent across all genotypes of a given comparison. Log2Fold Change (Log2FC) values were adjusted using DESeq2’s lfcShrink function with the ‘normal’ shrinkage estimator. Differentially expressed genes (DEGs) were defined as genes with FDR < 0.05 and absolute value of Log2FC > 0.2. All DEGs from bulk RNA-seq for all brain regions, ages and genotypes can be found in Table S1-S2.

DESeq2 was also used to run DE analysis between *Grin2a^+/-^* (4-week and 12-week) versus *Grin2b^+/C456Y^* 4-week samples for each brain region. For this analysis, all heterozygous and wild-type replicates for each region were read into a DESeq object. DESeq contrasts were then used to test for differences between the *Grin2a^+/-^* and *Grin2b^+/C456Y^*, taking into account the differences between batches. More formally, we considered the contrast defined by (*Grin2a^+/-^* - *Grin2a^+/+^*) - (*Grin2b^+/C456Y^* - *Grin2b^+/+^*). DEGs were defined as genes with FDR < 0.05 and absolute value of Log2FC > 0.2.

For comparing the transcriptomic changes in *Grin2a^+/-^* versus *Grin2a^-/-^*, the Pearson’s correlation r values were calculated between Log2FC values of the same brain regions in *Grin2a*^+/-^ versus *Grin2a*^-/-^. It should be noted that the use of the same set of wild types for *Grin2a^+/-^* and *Grin2a^-/-^* DE analysis may lead to over inflated correlation estimates.

A list of number of reads per replicate and alignment statistics for each brain region, genotype and age has been provided in Table S3. We did not notice a strong relationship between number of DEGs and average coverage, suggesting depth of sequencing is not the main driver of variation in number of DEGs between regions.

### snRNA-seq analysis

The Cell Ranger v3.0.2 pipeline (10x Genomics) (Zheng et al., 2017) was used to align reads from snRNA-seq to a mm10 mouse reference genome, which was built to include introns (https://cf.10xgenomics.com/supp/cell-exp/refdata-gex-mm10-2020-A.tar.gz). The -- chemistry=SC3Pv3 and --expect-cells=10000 flags were used in addition to default parameters for Cell Ranger count. Replicates for each brain region and age were downsampled to the same number of reads per cell using Cell Ranger aggr with default parameters. Ambient RNA removal was performed on the 12-week Hippocampus CellRanger aggr output, split by replicate, using CellBender v0.1.0, with the arguments --expected-cells 10000 and --total-droplets-included 25000 (Fleming et al., 2019). Quality control metrics for each sample used in this study may be found in Table S7.

Unique Molecular Identifier (UMI) counts were analyzed using the Seurat v3.2.3 (Stuart et al., 2019) package. Nuclei expressing less than 500 genes were removed and the remaining nuclei were normalized by total expression, multiplied by ten thousand, and log transformed. Seurat’s ScaleData function was then used to scale the data. Linear dimension reduction by PCA was performed using Seurat’s RunPCA function on the scaled data on variable genes, and significant PCs were identified. The nuclei were clustered in the PCA space using Seurat’s FindNeighbors and FindClusters functions then visualized using Uniform Manifold Approximation and Projection (UMAP) using significant PCs. Doublet identification was performed using the Scrublet v0.2.3 (Wolock et al., 2019) Python package with default parameters. Clusters and nuclei with a scrublet score above the region-specific thresholds were determined to be doublets and were removed from the data. The table below shows the used scrublet score thresholds for all snRNA-seq datasets:

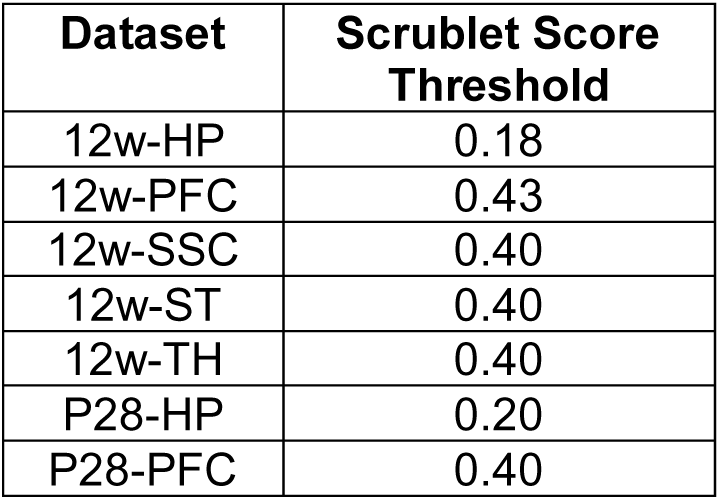

Cluster cell type identification was performed using the Azimuth R package (Hao et al., 2021). The references used for each brain region are summarized in the table below:

The references used for each brain region are summarized in the table below:
| Brain region | Azimuth Reference(s) Used |
| --- | --- |
| Hippocampus | Allen Brain Atlas "WHOLE CORTEX & HIPPOCAMPUS – 10X GENOMICS" (Yao et al., 2021b), DropViz<br>"annotation.BrainCellAtlas_Saunders_version_2018.04.01.rds"<br>(Saunders et al., 2018) |
| PFC | Azimuth Mouse Motor Cortex(Yao et al., 2021a), Allen Brain Atlas<br>"WHOLE CORTEX & HIPPOCAMPUS – 10X GENOMICS" |
| SSC | Azimuth Mouse Motor Cortex |
| Striatum | Allen Brain Atlas "WHOLE CORTEX & HIPPOCAMPUS – 10X GENOMICS", DropViz<br>"annotation.BrainCellAtlas_Saunders_version_2018.04.01.rds" |
| Thalamus | Allen Brain Atlas "WHOLE CORTEX & HIPPOCAMPUS – 10X GENOMICS", DropViz<br>"annotation.BrainCellAtlas_Saunders_version_2018.04.01.rds" |

These cell type assignments were then confirmed using the region-specific cell type marker genes. Neuronal subtypes were similarly identified by re-clustering nuclei of the given major cell type and confirming the Azimuth subtype identity labels through marker gene expression. Marker genes for the more widely classified major cell types and subtypes were identified in the literature (Ding et al., 2020). Seurat’s FindMarkers function was used to identify marker genes for rarer cell types and subtypes. These markers were then queried on mouseBrain.org (Zeisel et al., 2018) and or dropviz.org (Saunders et al., 2018) to evaluate the cell type identity. A full list of the marker genes used in this study may be found in Fig. S2F-M and in Table S7. A list of number of nuclei per cell type per sample can be found in Tables S22-28.

The propeller function from the speckle R v0.0.3 package (Phipson et al., 2022) was used with arcsin normalization to test if there were statistical differences in major cell type and subtype composition across genotypes within each dataset.

### Pseudocell differential expression analysis of snRNA-seq

For differential expression analysis, we employed a pseudocell approach for each age and brain region that aggregates expression of every 40 nuclei with similar transcriptomes within each replicate, cell type, and subtype. The pseudocell method resolves known technical variation issues of snRNA-seq transcriptome data including dropout expressions and pseudoreplication through averaging expression of similar nuclei (Cao et al., 2018; Kanton et al., 2019). To construct pseudocells, single nuclei were first grouped based on replicate and cell type. Within each group, pseudocell centers were identified by applying k-means clustering on the top 30 principal components (PCs) with k set so that each cluster contains 20 nuclei at minimum and 40 nuclei on average. We also weighted PCs by their variance explained to ensure the stability of the results across different group sizes (i.e., cell types per replicate). Random walk methods are shown to have superior performance in identifying nuclei with similar transcriptome profiles compared with the spherical, distance-based methods (Stuart et al., 2019; van Dijk et al., 2018). Therefore, we assigned nuclei to the pseudocell centers (i.e., k-means centroids) using a random walk approach on the cell-cell distances in the PCA space (van Dijk et al., 2018). Lastly, we constructed pseudocell samples by aggregating the raw UMI counts of nuclei assigned to each pseudocell. This resulted in representation of each cell type or subtype from each replicate by multiple pseudocells. Cell types and subtypes that were identified but did not have enough nuclei present to create at least two pseudocells per replicate were removed. Differential expression analysis was performed using limma’s duplicateCorrelation mixed model analysis function with robust empirical Bayes moderated t-statistics and mouse ID as random effect (Law et al., 2014). Genotype and the percentage of mitochondrial reads were used as covariates for all analyses. Experiment batch number was added as a covariate for *Grin2a* 12-week hippocampus (12w-HP) and Grin2a 12-week PFC, where samples were sourced from two separate sequencing runs. Samples of each batch are indicated in Table S23 for PFC 12-week and in Table S25 for HP 12- week.

Genes with Benjamini-Hochberg adjusted p-value of less than 0.05 and absolute Log2FC > 0.2 were deemed as DEGs. To further ensure the robustness of the identified DE genes, we calculated the log fold-change pattern (summarized as up or down regulated) of every pairwise combination of experiment (either HT or KO samples) versus WT samples. For example, a total of 3 WT samples x 3 KO samples = 9 comparisons for KO and 3 WT samples x 4 HT samples = 12 comparisons for HT DE genes. As a measure of robustness, we next calculated the fraction of comparisons in which the log fold-change had a similar direction to that of the mixed linear model. We observed this measure of robustness is above 0.8 for the majority (∼ 60%) of the DEGs, indicating that the fold-change pattern of DEGs is shared across samples rather than being driven by a subset. We retained DEGs with robustness score above 0.8 as DEGs. A list of all DEGs for all cell types, brain regions and ages from pseudocell analysis can be found in Tables S22-28.

The clusters that were identified in the initial clustering for each brain region but removed for the pseudocell analysis due to low nuclei counts are as follows: **Hippocampus (4- and 12-week)**: Cajal Retzius, Endothelial, Ependyma, SST Chodl inhibitory subtype; **PFC (4- and 12-week) and SSC (12-week)**: Endothelial, Vascular, SST Chodl inhibitory subtype, L6b excitatory subtype (additionally, L5/6 NP excitatory subtype was removed from 12-week SSC); **Striatum (12-week)**: Choroid Plexus, Endothelial, Vascular, the cholinergic Interneuron_Chat subtype; **Thalamus (12- week)**: Endothelial, Ependyma, Vascular, the DEINH3 and MEINH11 inhibitory subtypes.

### Pseudobulk differential expression analysis of snRNA-seq

We also utilized a pseudobulk approach (Lun and Marioni, 2017) for differential expression analysis. For this analysis, counts were grouped by cell type and replicate. All gene counts were summed across each replicate within each cell type, resulting in one count value per replicate per genotype. Differential expression analysis was then run using the EdgeR v3.22.3 R package (Robinson et al., 2010). Experiment batch number was added as a covariate for *Grin2a* 12w-HP and *Grin2a* 12w-PFC, where samples were sourced from two separate sequencing runs. Genes with Benjamini-Hochberg adjusted p-value of less than 0.05 were deemed as DEGs. A list of all DEGs for all cell types, brain regions and ages from EdgeR analysis can be found in Tables S8- 14.

### Gene set enrichment analysis

The DESeq2 t-stat values from bulk RNA-seq analysis or the Log2FC values from pseudocell or EdgeR analysis were extracted to run Gene Set Enrichment Analysis (GSEA) (Subramanian et al., 2005) using the fGSEA v1.16.0 package (Korotkevich et al., 2021) and either the C5 v7.2 gene set collection (including 14,765 Gene Ontology (GO) terms) from Molecular Signature Database (http://www.gsea-msigdb.org/gsea/msigdb), SynGO collection (Koopmans et al., 2019) or gene sets generated from the literature. For proteomics data, proteins were pre-ranked using their moderated t-statistic within each comparison (*Grin2a^+/-^* vs wild type or *Grin2a^-/-^* vs wild type). For proteins with multiple isoforms, the isoform with the largest number of spectra was used for pre-ranking. For each of these gene sets, mouse gene symbols from transcriptomics and proteomics results were mapped to matching *Homo sapiens* homolog-associated gene symbols through annotations extracted from Ensembl’s BioMart data service; the human symbols were then used to perform GSEA. A list of all significant GO terms (FDR < 0.05) for all datasets are provided in the following tables: Bulk RNA-seq (Tables S4-5); snRNA-seq EdgeR analysis (Tables S15-21); snRNA-seq pseudocell analysis (Tables S29-35); proteomics (Table S37).

For the ‘*activity regulated’* gene-set, the neuronal activity-induced rapid, delayed and slow primary response genes defined by (Tyssowski et al., 2018) were compiled. For the ‘*Dopamine-induced*’ gene-set, significantly (FDR < 0.1) upregulated DEGs in striatal neuronal culture following one hour dopamine treatment (Savell et al., 2020) were used. The “*cAMP-induced CREB target*’ gene- set were identified by (Zhang et al., 2005). For a list of curated gene-sets please see Table S6.

### Quantitative Mass Spectrometry analysis

The data was searched on Spectrum Mill MS Proteomics Software (Broad Institute) using mouse database that contained 47,069 entrees downloaded from Uniprot.org on 12/28/2017. The Spectrum Mill generated proteome level export which was filtered for proteins identified by two or more peptides for further analysis. Protein quantification was achieved by taking the ratio of TMT reporter ions for each sample the median of all channels. TMT16 reporter ion intensities were corrected for isotopic impurities in the Spectrum Mill protein/peptide summary module using the afRICA correction method which implements determinant calculations according to Cramer’s Rule and correction factors obtained from the reagent manufacturer’s certificate of analysis (https://www.thermofisher.com/order/catalog/product/90406) for lot number VA299611.

After performing median-MAD normalization, a moderated two-sample t-test was applied to the datasets to compare wild-type, *Grin2a*^+/-^ and *Grin2a*^-/-^ sample groups. Significant differentially expressed proteins (DEPs) were defined as proteins with nominal p-value < 0.05 and absolute value of Log2FC > 0.2. A list of all DEPs for *Grin2a*^+/-^ and *Grin2a*^-/-^ hippocampal samples are provided in Table S36.

### EEG analysis

Sleep state classification (NREM, REM or Wake) was performed on 10 second epochs of EEG/EMG data using a machine learning model as described previously (Herzog et al., 2022). LUNA software for EEG analysis (http://zzz.bwh.harvard.edu/luna) was used to compute absolute power for each oscillatory band (slow: 0.5-1 Hz, delta: 1-4 Hz, theta: 4-8 Hz, alpha: 8-12 Hz, sigma: 12-15 Hz, beta: 15-30 Hz, gamma: 30-50 Hz) for NREM sleep during the light cycle, where mice are predominantly asleep. Statistical differences between wild type, *Grin2a^+/-^* and *Grin2a^-/-^* animals were computed using one-way ANOVAs with post hoc Tukey-Kramer tests for multiple comparisons using MATLAB (MathWorks, Natick, MA, RRID: SCR_001622).

### MoSeq analysis

The depth videos were processed in the extraction pipeline as described previously (Wiltschko et al., 2020). The kinematic values such as velocity, angle, the position of the mouse’s centroid, position to center, length, height, and width were extracted from the videos. The depth images were cropped to 80 pixels by 80 pixels and processed such that the mouse center was at the frame center and the nose was always pointing right. The extracted frames went through principal component analysis to reduce the dimensionality and the top 10 principal components (PCs) were input into the MoSeq autoregressive hierarchical Dirichlet process hidden Markov model (AR- HMM) to learn the behavioral motifs, defined as syllables. The model was trained in robust mode, such that the noise of the hidden states in AR-HMM were sampled from a t-distribution. Model hyperparameter, kappa, which was the self-transition bias parameter that controlled the average syllable durations was selected after scanning through a range of kappa values. The syllable duration distributions in the models trained with different kappa values were compared with the block duration distributions from a model-free changepoint detection model. The kappa value that yielded a syllable duration median that best matched that of the model-free block duration was selected for training the final models. The final models with the kappa found in kappa scan (kappa = 1149215) trained with 1,000 iterations of Gibbs sampling and the model that had the median log likelihood was selected for further data analysis. The final single model included all 16 wildtype, 9 *Grin2a*^+/-^ and 13 *Grin2a*^-/-^ mice in the MoSeq open field assay recordings. The model fitting yielded 49 syllables that explains 99% of the total frames in the session.

Two sessions were filtered out in the behavioral analysis due to abnormal mouse sizes. The behavioral summaries of an example wild-type mouse was computed from a 5 min segment in the middle of the example session. The speed was calculated as the absolute 2D distance traveled between two frames divided by 1/30 second (sampling frame rate 30 fps). The length was computed as the length of the mouse body contour in each frame. The distance to center was computed as the magnitude of the centroid of the animal to the center of the detected region of interest (ROI) that represented the arena, normalized by the radius of the ROI.

Syllable usages are computed from syllable orders within each session and normalize within each group. Differences between the mean velocity and distance traveled between two groups are done with Welch’s t-test from scipy.stats.ttest_ind with equal_var=False, and permutation test. In the permutation test, the number of permutations is 99999 and 1 pseudo count is added in computing p value.

For the clozapine (10mg/kg) or haloperidol (0.25 mg/kg) analysis, we used publicly available MoSeq model syllable label results at https://github.com/dattalab/moseq-drugs. The modeling procedures were described in detail in (Wiltschko et al., 2020) and the data was recorded using Kinect for Windows v.1 instead of Kinect for Windows v.2 in this experiment.

## DATA AVAILABILITY

RNA-seq and snRNA-seq data that support the findings of this study will be deposited at Gene Expression Omnibus (GEO) and at the Broad Single Cell Portal (scPortal). The GEO accession number and scPortal link will be provided.

The original mass spectra and the protein sequence databases used for searches have been deposited in the public proteomics repository MassIVE (http://massive.ucsd.edu) and are accessible at ftp://MSV000090487@massive.ucsd.edu with the password: astrocytes. If requested, also provide the username: MSV000090487. These datasets will be made public upon acceptance of the manuscript.

## CODE AVAILABILITY

The code used for RNA-seq and snRNA-seq analysis using EdgeR is available on GitHub at https://github.com/aanicolella/grin2a. The code for snRNA-seq analysis using pseudocell will be made public upon acceptance of the manuscript. The MoSeq model syllable label results used in this study may be found on GitHub at https://github.com/dattalab/moseq-drugs.

## ACKNOWLEDGMENTS

We thank Charles Vanderburg for assisting with mouse brain dissection, and Naeem N. Nadaf, Nicolas Lapique and Xian Adiconis for assistance with sequencing library preparations. We thank Matthew Beck and Tyler Caron for assisting with maintaining the *Grin2a* mouse colony. We thank Kevin Mastro, Gordon Fishell and Prabhat Kunwar for invaluable feedback. Research reported in this manuscript was supported by the Stanley Center for Psychiatric Research. Z.F. is supported by an Otto Hahn Fellowship of the Max Planck Society. M.S. serves on the Scientific Advisory Board of Biogen, Cerevel, Neumora, Vanqua Bio, and ArcLight Therapeutics.

## AUTHOR CONTRIBUTIONS

Z.F. and M.S. designed the experiments. Z.F. performed all brain dissections and sequencing library preparation for single-nucleus and bulk RNA-seq experiments with assistance from K.S.B. N.S. and K.S.B. performed animal perfusion and brain collection of *Grin2a* mutant mice. N.S. performed MoSeq experiments and Western blotting. K.S.B. performed qRT-PCR. A.N., S.S. and Z.F. performed RNA-seq analysis with critical contributions by J.L. B.S. assisted with RNA-seq analysis and data interpretation and visualization. V.G. and E.M. developed and provided the pseudocell algorithm. B.D. performed PSD fractionation and supervised the proteomics analysis and interpretation. K.B. performed proteomics experiments supervised by H.K. and S.A.C. S.A. performed proteomics data analysis, PSD fractionation for Western blot validation and figure preparation for the proteomics result. S.L. performed MoSeq analysis and figure preparation for the MoSeq result supervised by S.R.D. L.H. performed EEG experiment and analysis. W.S. performed animal perfusion and brain collection of *Grin2b* mutant mice. E.K. assisted with *Grin2b* data analysis. M.S. conceived and supervised all aspects of the project. Z.F., A.N. and M.S. wrote the manuscript with input from all authors.

